# *Tet2*-Clonal Hematopoiesis Promotes Whereas *Dnmt3a*-CH Inhibits Breast Cancer Progression During Aging

**DOI:** 10.64898/2026.11.09.765294

**Authors:** Nagihan G. Gonullu, Karolina A. Zielińska, Jonas Fullin, Francesco Prisco, Steffen Boettcher, Francisco Caiado, Markus G. Manz

**Author notes:** Authors contributed equally (Francisco Caiado, Markus G. Manz).

## Abstract

Clonal hematopoiesis of indeterminate potential (CHIP) driven by somatic mutations in blood cells increases with age and is associated with adverse outcomes in cancer patients. How CHIP mutations influence solid tumor progression remains unclear. Using inducible murine models of *Dnmt3a*-or *Tet2*-mutant CHIP combined with orthotopic syngeneic breast cancer models, we demonstrate that upon aging *Dnmt3a*-mutant CHIP prevents whereas *Tet2*-mutant CHIP drives breast cancer progression. *Dnmt3a*-mutant immune infiltrates established interferon-mediated MHCII^high^ tumor-associated macrophages (TAMs) that supported cytotoxic CD8+ T cell-mediated tumor rejection. In contrast, *Tet2*-mutant TAMs exhibited an inflammatory, immunosuppressive phenotype characterized by increased IL-1 and CXCL2 production, reduced MHCII expression, and impaired T cell recruitment and activation. Therapeutic blockade of IL-1R1 or CXCR2 reduced tumor progression in aged *Tet2*-mutant CHIP mice and partially restored anti-tumor immunity. These findings establish that the effects of CHIP on cancer are mutation- and age-dependent and identify IL-1/CXCL2-driven inflammation as a therapeutic vulnerability of tumor-promoting *Tet2*-mutant CHIP.

## INTRODUCTION

Accumulation of somatic mutations and subsequent clonal selection for fitness-enhancing mutations are hallmarks of aging. In the hematopoietic system, this process can give rise to Age-Related Clonal Hematopoiesis (ARCH), also termed Clonal Hematopoiesis of Indeterminate Potential (CHIP). CHIP is a form of CH defined by the presence of one or more somatic mutations in known myeloid leukemia driver genes (most frequently *DNMT3A*, *TET2* and *ASXL1*), detected in hematopoietic cells at a variant allele frequency (VAF) of at least 2% in individuals without an overt hematological malignancy or unexplained cytopenia^1,2^. CHIP incidence increases with age and is associated with increased risk of blood cancer, cardiovascular and inflammatory diseases, infections and all-cause mortality^3,4^.

Recent findings suggest that CHIP is also associated with increased solid cancer incidence and growth. Accordingly, CHIP carriers with larger clones (VAF over 10%) have increased risks of lung, skin, prostate and breast cancer^5^, while CHIP in cancer patients is associated with increased mortality from disease progression^6,7^. Recently, tumor-infiltrating CH (TI-CH), defined by CHIP mutations at a VAF of at least 2% in at least one tumor region, has emerged as an important prognostic factor and is associated with shorter recurrence-free survival and increased mortality compared with CHIP confined to blood^8^. Moreover, tumors from TI-CH cases show enrichment for myeloid immune and inflammatory signatures, particularly IL-1-dependent transcriptomic signatures in aged individuals^9^. Importantly, loss-of-function mutations in *DNMT3A* and *TET2* (accounting together for over 75% of all CHIP mutations) can induce hyperinflammatory responses in mutant mature myeloid cells, potentially contributing to CHIP-associated health risks^7^. However, how CHIP and its ensuing inflammation contribute to solid tumor development and outcome has not been firmly established. Studies of hematopoietic *Tet2* loss have reported divergent effects across tumor models^10,11^, suggesting that its impact may depend on the specific mutation, hematopoietic cell population carrying the mutation, tumor type, and potentially environmental context such as age^10^.

Breast cancer ranks first in incidence and mortality among women. CHIP is associated with increased incidence of breast cancer^5,12^ and is frequent in breast cancer patients^6^. Moreover, immune cells belonging to the CHIP clone have been detected in the TME of breast cancer patients^13^. CHIP has additionally been linked to increased chemotherapy-related hematological and cardiac toxicities in breast cancer patients^14,15^. However, the impact of CHIP on breast cancer progression and on patient survival remains unclear, with studies reporting either no association^16–18^ or disease-promoting effects^19,20^. Importantly, the direct influence of CHIP on breast cancer, immune cell composition and function remains unknown.

Here, we investigated the effects of *Tet2*- and *Dnmt3a*-mutant CHIP on breast cancer progression using inducible transgenic mouse models of CHIP^21^ combined with orthotopic syngeneic breast cancer models. We further examined how these mutations shape the tumor immune microenvironment and explored potential therapeutic implications.

## RESULTS

### Breast cancer progression depends on the CHIP mutation profile in young mice

To investigate the effects of the most frequent forms of CHIP, *Tet2*- and *Dnmt3a*-mutant CHIP, on breast tumor growth and metastasis, we used our previously established triple transgenic hematopoietic-specific, tamoxifen-inducible mouse models of *Tet2* loss and *Dnmt3a* R878H mutation, both carrying a *ZsGreen* reporter^21^. Tamoxifen administration induced either *Tet2* heterozygosity or the *Dnmt3a* R878H mutation together with *ZsGreen* expression, enabling tracing of mutant hematopoietic cells. Mice carrying wild-type (WT) *Tet2* or *Dnmt3a* alleles served as controls. To assess tumor progression, we established a syngeneic orthotopic breast cancer model using E0771 cells, transduced with a dual-reporter expression vector. (**Figure S1A**). To induce CHIP, young (2–4-month-old) triple-transgenic mice received a single high dose of tamoxifen (50 mg/kg), resulting in a high blood clonal fraction (∼70% ZsG⁺ leukocytes) (**Figure S1B**). Following E0771 transplantation and longitudinal tumor burden quantification, there was no impact on primary tumor growth in young *Tet2*^+/-^-CHIP compared with WT mice (**Figure S1C-F**). Quantification of metastatic tumor spreading by ex vivo imaging of multiple organs revealed that *Tet2*^+/-^-CHIP associates with increased lung metastasis compared to controls (**Figure S1G-H**). In contrast, young *Dnmt3a^R^*^878^*^H/+^*-CHIP mice showed both reduced primary tumor growth (**Figure S1J-M**) and lung matastasis compared to WT (**Figure S1N-P**). Thus, in young mice, *Dnmt3a*-mutant CHIP suppresses breast cancer progression, whereas *Tet2*-mutant CHIP primarily increases metastatic burden.

### *Tet2^+/-^*-CHIP enhances whereas *Dnmt3a^R^*^878^*^H/+^*-CHIP inhibits breast cancer progression in middle-aged mice

As both CHIP and breast cancer are more frequent in the elderly population^17^, we evaluated how *Tet2*- and *Dnmt3a*-mutant CHIP affects breast cancer progression in aging mice. We aged mice carrying high *Tet2-* or *Dnmt3a-*mutant clonal fractions (70-90% ZsG+ leukocytes) to 10-14 months (termed middle-aged) and transplanted them orthotopically with E0771 breast cancer cells (**Figure 1A-B**). Strikingly, middle-aged mice with *Tet2*^+/-^-CHIP showed significantly increased breast tumor growth, assessed by longitudinal total flux quantification, tumor volume and terminal tumor weight (**Figure 1C-F**). Moreover, *ex vivo* and image analysis of histological sections of lungs revealed enhanced metastatic dissemination (**Figure 1G-J**). In contrast, primary tumor growth was reduced in *Dnmt3a^R^*^878^*^H/+^*-CHIP middle-aged mice, while lung metastatic burden was comparable to controls (**Figure 1 K-R**). These results suggest that age influences breast cancer progression in case of mice carrying high clonal sizes of *Tet2^+/-^-*CHIP, while its impact in WT and *Dnmt3a^R^*^878^*^H/+^*-CHIP mice is minimal (**Figure S2A-E**). Using the non-metastatic breast cancer cell line Py230 in the syngeneic orthotopic model, we confirmed the tumor-promoting effect of *Tet2^+/-^-*CHIP in middle-aged mice (**Figure S3A-F**). To understand if the tumor-promoting effect of *Tet2^+/-^-*CHIP depends on mutant clonal size, we repeated the experiment in middle-aged mice induced with a lower *Tet2*-mutant clonal fraction (25-35% ZsG+ leukocytes) (**Figure S4A-B**). Even at lower clonal sizes, *Tet2^+/-^-*CHIP enhanced breast cancer growth, while lung metastasis increased numerically but did not reach statistical significance (**Figure S4C-H**). However, the effect on primary tumor growth was smaller than in mice carrying higher *Tet2*-mutant clonal sizes (**Figure S4I**), confirming clonal size as a relevant determinant of the pro-tumoral effect of *Tet2*-mutant CHIP, as previously shown for other CHIP-associated pathologies^4^. Overall, these findings demonstrate that in middle-aged mice, breast tumor progression is influenced by CHIP in a mutation and clone size-dependent manner, with *Tet2^+/-^-*CHIP promoting and *Dnmt3a^R^*^878^*^H/+^*-CHIP inhibiting breast cancer progression.

**Figure 1:**
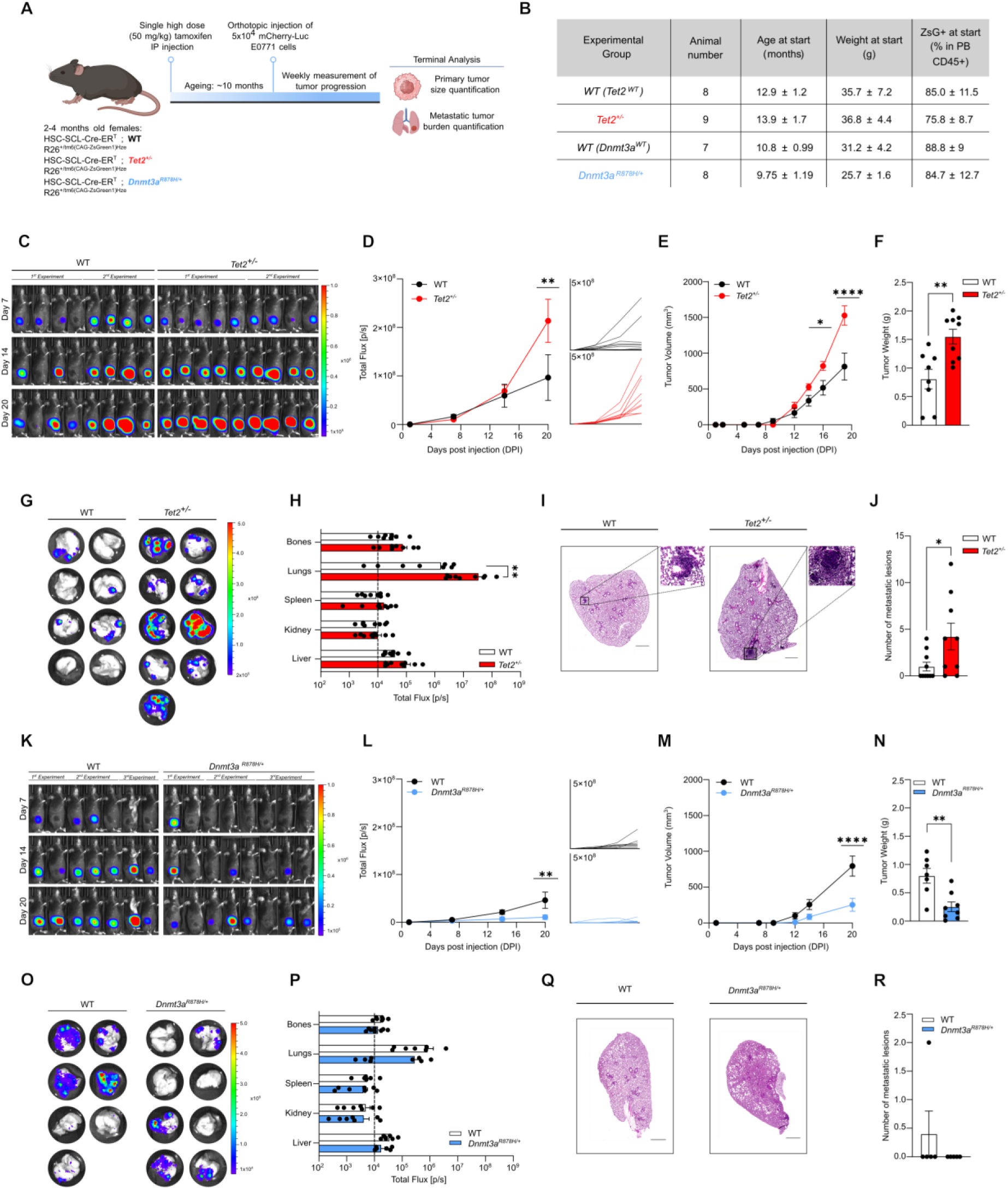
*Tet2*-mutant CHIP promotes while *Dnmt3a*-mutant CHIP reduces breast cancer progression in middle-aged mice. (A) Graph depicting experimental outline. (B) Graph depicting animal numbers, age, weight and peripheral blood chimerism (CD45+ ZsG+) of each experimental group. (C) *In vivo* growth of orthotopic injected, mCherry-Luciferase expressing E0771 breast cancer cells over time, evaluated by IVIS imaging of WT and *Tet2^+/-^*CHIP animals. (D) Tumor growth curves (left panel) and individual curves (right panel) determined by quantification of luminescent intensity from images in (C). (E) Tumor volume measurements over time in mm^3^. (F) Tumor weight measurement in grams upon termination of experiment on day 21. (G) *Ex vivo* IVIS images of lungs showing luciferase expressing metastasis in WT and *Tet2^+/-^*CHIP animals. (H) *Ex vivo* quantification of luminescent intensity in indicated organs of WT and *Tet2^+/-^*animals. Dashed line represents the average luminescent background signal (I) Representative Hematoxylin and Eosin (HE) stained lungs section, scale bar 800µm with enlargement of indicated areas. (J) Quantification of numbers of metastases in the lungs from HE staining. (K) *In vivo* growth of orthotopic injected, mCherry-Luciferase expressing E0771 breast cancer cells over time, evaluated by IVIS imaging of WT and *Dnmt3a^R^*^878^*^H/+^* CHIP animals. (L) Tumor growth curves (left panel) and individual curves (right panel) determined by quantification of luminescent intensity from images in (K). (M) Tumor volume measurements over time in mm^3^. (N) Tumor weight measurement in grams upon termination of experiment at day 21. (O) *Ex vivo* IVIS images of lungs showing luciferase expressing metastasis from WT and *Dnmt3a^R^*^878^*^H/+^* CHIP animals. (P) *Ex vivo* quantification of luminescent intensity in indicated organs of WT and *Dnmt3a^R^*^878^*^H/+^* CHIP animals. Dashed line represents the average luminescent background signal. (Q) Representative HE stained lungs section, scale bar 800µm with enlargement of indicated areas. (R) Quantification of numbers of metastases in the lungs from HE staining. For panels (C-J), data is pooled from two independent experiments and for panels (K-P) data is pooled from three independent experiments. Statistical analysis was performed by using two-way ANOVA (D, E, H, L, M, P) and unpaired t test (F, J, N, R). Data is shown as mean with SEM. *p<0.05, **p<0.01, ***p<0.001, ****p<0.0001.

### Tumor cells from middle-aged mice with *Tet2^+/-^-*CHIP downregulate interferon response and increase NF-kB pathway transcriptomic signatures

To investigate how *Tet2-* and *Dnmt3a*-mutant CHIP affect tumor growth, we performed bulk RNA sequencing on sorted mCherry+ E0771 tumor cells from WT, *Tet2^+/-^* or *Dnmt3a^R^*^878^*^H/+^*-CHIP middle-aged mice (**Figure 2A**). Principal component analysis revealed distinct transcriptional profiles among tumors from mice carrying different CHIP mutations (**Figure 2B**). Next, we identified differentially expressed genes (DEGs, **Table S1**) between tumor cells from *Tet2^+/-^*over WT, *Dnmt3a^R^*^878^*^H/+^* over WT or *Tet2^+/-^*over *Dnmt3a^R^*^878^*^H/+^*CHIP animals (**Figure 2C**) and performed gene set enrichment analysis (GSEA) with the Hallmark collection. Compared with both WT and *Dnmt3a^R^*^878^*^H/+^*-CHIP mice, tumors from *Tet2*^+/−^-CHIP mice showed downregulation of IFNα and IFNγ response signatures and upregulation of epithelial-to-mesenchymal transition and NF-κB signaling (**Figure 2D**). The Hallmark IFNγ response signature indicated reduced adaptive immune activity in tumors from *Tet2*^+/−^-CHIP mice, including downregulation of *Tap1* and *Tapbp* involved in Major Histocompatibility Complex I (MHCI), and *H2-Aa*, *H2-DMa*, *Ciita*, and *Cd74* involved in MHCII. IFNγ-responsive T-cell chemoattractants *Cxcl9* and *Cxcl10* were also reduced (**Figure 2E, upper panel**). Consistently, the Hallmark Allograft Rejection pathway showed downregulation of *B2m*, *Tap1*, *Tap2*, *Cxcl9*, *H2-Aa* and *H2-DMa* in tumors from *Tet2^+/-^*-CHIP mice (**Figure 2E, lower left panel**). Conversely, expression of NFkB-induced genes, such as *Nfkbia, Tnfaip2/3/6* and *Nr4a1* increased in tumors from *Tet2^+/-^*-CHIP mice compared to WT and *Dnmt3a^R^*^878^*^H/+^*-CHIP mice (**Figure 2E lower right panel**). Overall, these analyses revealed distinct transcriptional profiles associated with *Tet2*- and *Dnmt3a*-mutant CHIP. Enhanced tumor growth in middle-aged *Tet2*^+/−^-CHIP mice were associated with increased NF-κB-associated innate inflammatory signaling and reduced interferon-responsive adaptive immune signaling, resulting in decreased MHC expression and potentially impaired anti-tumor immunity.

**Figure 2:**
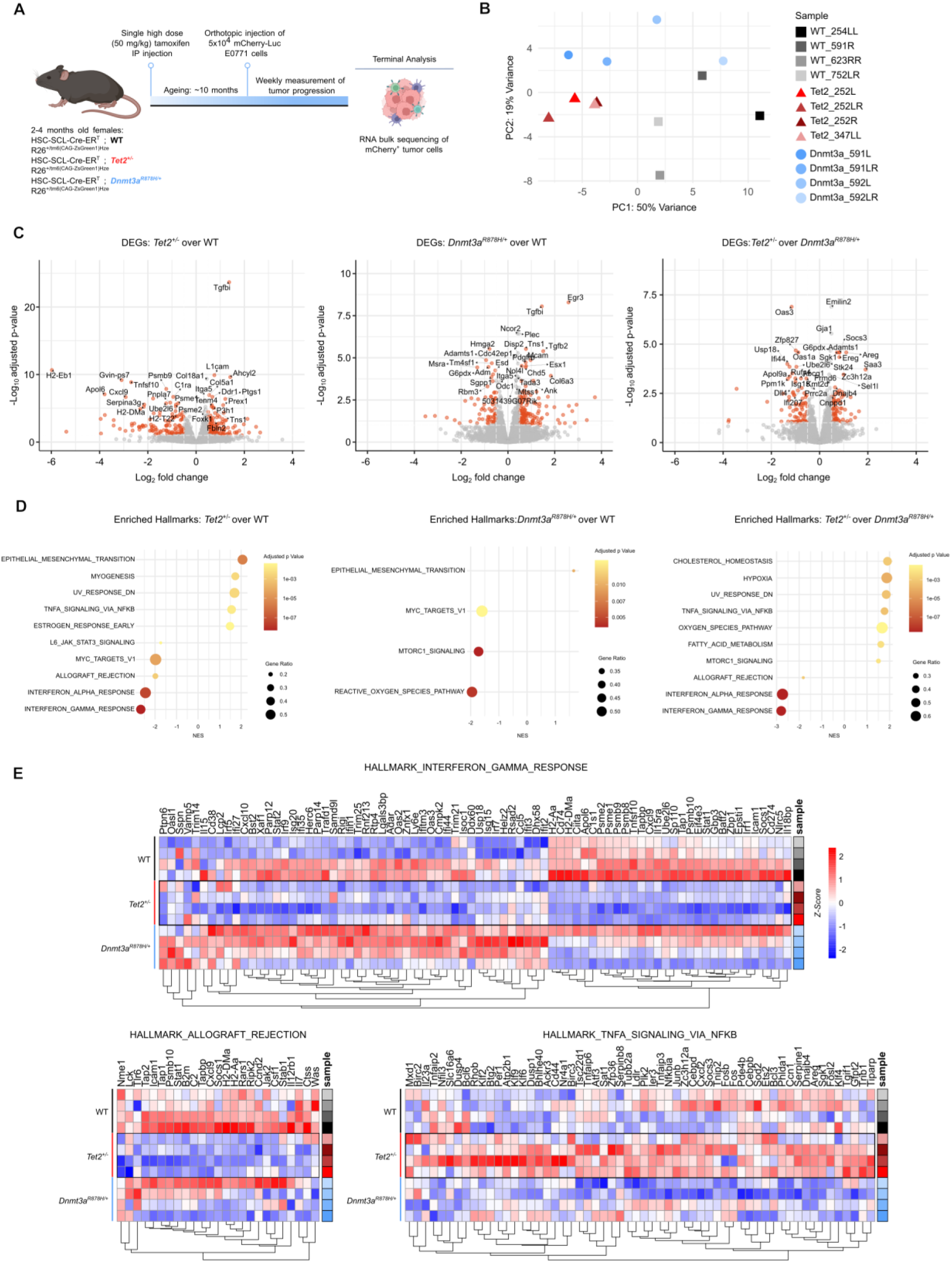
Breast cancers from middle-aged *Tet2*-mutant CHIP mice show increased NFkB and diminished IFNα/γ response signatures compared to WT and *Dnmt3a*-mutant CHIP mice. (A) Graph depicting experimental outline. (B) Principal component (PC) analysis of bulk RNA sequencing of breast cancers from WT, *Tet2^+/-^* and *Dnmt3a^R^*^878^*^H/+^* CHIP animals. (C) Volcano plots showing differentially expressed genes (DEGs, -1 > Log2 fold change > 1; Adjusted p-value < 0.05) comparing cancers from *Tet2^+/-^*versus WT, *Dnmt3a^R^*^878^*^H/+^* versus WT and *Tet2^+/-^*versus *Dnmt3a^R^*^878^*^H/+^* genotypes. (D) Gene set enrichment analysis of cancer cells showing enriched hallmark datasets from *Tet2^+/-^*vs WT, *Dnmt3a^R^*^878^*^H/+^*vs WT and *Tet2^+/-^*vs *Dnmt3a ^fl-R^*^878^*^H/+^* genotypes. (E) Heatmap showing the expression of genes in datasets Hallmark Interferon Gamma Response (upper panel), Allograft Rejection (lower left panel) and TNFα Signaling Via NFkB (lower right panel) distributed by WT, *Tet2^+/-^* and *Dnmt3a ^fl-R^*^878^*^H/+^* CHIP background from which cancers were extracted.

### *Dnmt3a^R^*^878^*^H/+^* and *Tet2^+/-^*-CHIP result in divergent tumor immune compositions with *Tet2*-mutant cells promoting immunosuppression in middle-aged mice

To uncover immunological alterations in tumor-bearing middle-aged WT, *Tet2^+/-^*or *Dnmt3a^R^*^878^*^H/+^*-CHIP mice, we characterized the immune composition of peripheral blood (PB), bone marrow (BM) and primary E0771 tumors by flow cytometry (**Figure 3A, Figure S5, Figure S6**). Tumor inoculation induced CD11b+ myeloid expansion and B/T-cell contraction in PB and BM across genotypes, consistent with cancer-adapted hematopoiesis^22^, but these changes were more pronounced in *Tet2^+/-^*^-^CHIP mice (**Figure 3B, Figure S5A**). Tumor-infiltrating CD45+ immune cells negatively correlated with tumor weight, with *Tet2^+/-^*-CHIP-bearing tumors showing lower immune infiltration and higher tumor weight, whereas *Dnmt3a^R^*^878^*^H/+^*-CHIP-bearing tumors showed the opposite pattern (**Figure 3C-D**). The tumor immune composition differed drastically between genotypes. Consistently, *Tet2^+/-^*-CHIP-bearing tumors were enriched in neutrophils and macrophages (representing over 50% of the TME) and depleted of T cells (under 25%), particularly CD8+ T cells, whereas *Dnmt3a^R^*^878^*^H/+^*-CHIP-bearing tumors showed the opposite pattern, with fewer neutrophils/macrophages and increased CD4+, CD8+, and CD4-CD8-(DN) T cells (**Figure 3E-F**). Reduced CD4+ and CD8+ T-cell frequencies in *Tet2^+/-^*-CHIP-bearing tumors persisted after normalization to tumor weight, supporting an immunosuppressive TME (**Figure 3G**). These immune alterations were also partially detected in Py230 tumors from middle-aged *Tet2^+/-^*-CHIP mice **(Figure S3G-N)** and in mice with lower *Tet2*-mutant clonal fractions (**Figure S4J-Q**), but were absent in young mice (**Figure S7A-G**).

**Figure 3:**
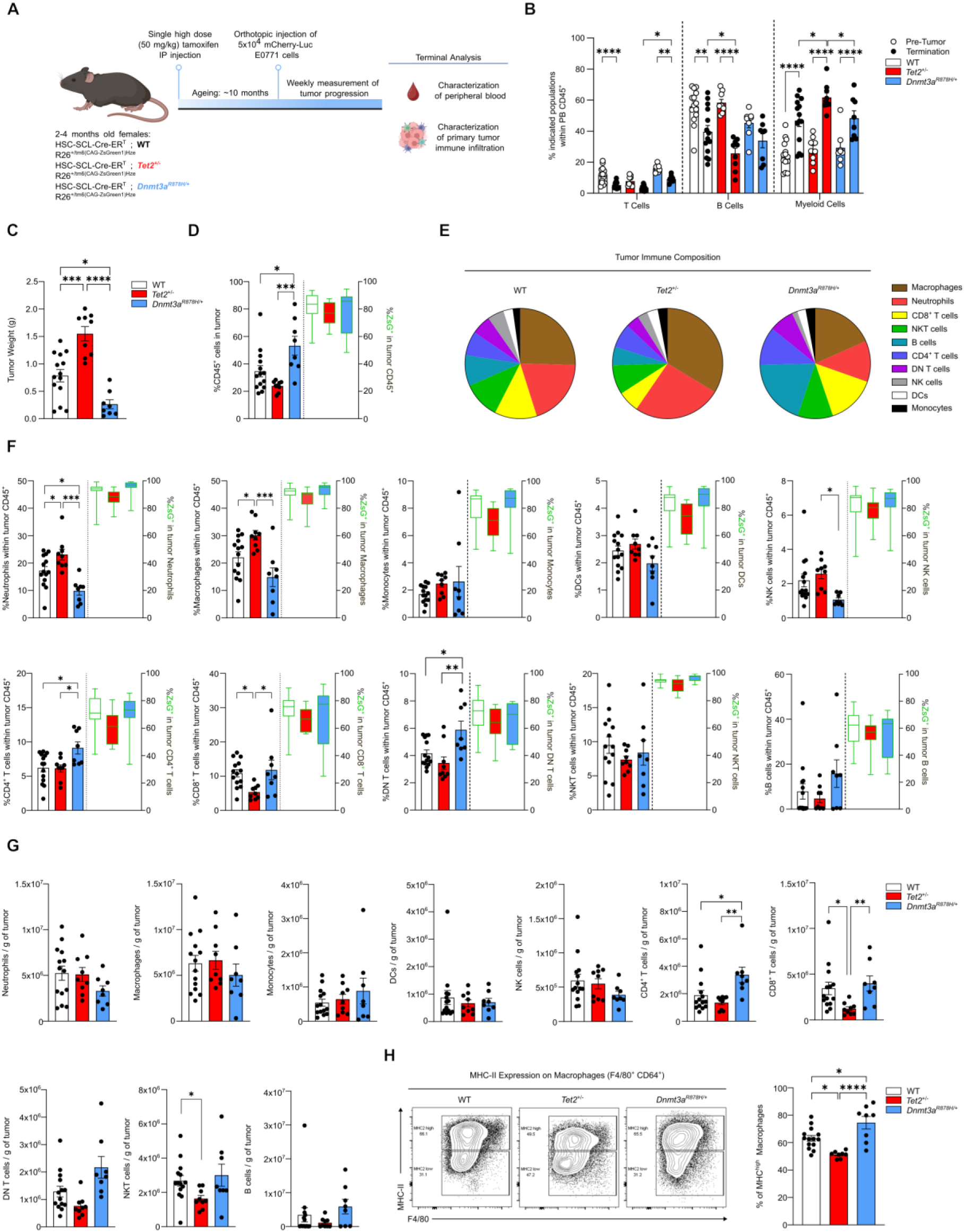
Breast cancers from middle-aged mice with *Tet2*-mutant CHIP contain increased macrophages and neutrophils and reduced CD8+ T cell infiltration compared to WT and *Dnmt3a*-mutant CHIP mice. (A) Graph depicting experimental outline. (B) Frequency of T cells, B cells and myeloid cells (CD11b+) in peripheral blood (PB) within CD45+ leukocytes before orthotopic breast cancer injection and at terminal analysis. (C) Tumor weight in grams at terminal analysis (g). Data is pooled from the groups shown in Figure 1 F and N. (D) Percentage of total CD45+ immune cells in total live cells of tumor microenvironment in WT, *Tet2^+/-^* and *Dnmt3a^R^*^878^*^H/+^* CHIP animals by flow cytometric analysis. Frequency of ZsG+ mutant immune cells in the indicated population are depicted on the right y-axis. (E) Pie chart depicting tumor immune composition as a percentage of indicated immune populations in CD45+ leukocytes. (F) Flow cytometric analysis of indicated tumor-infiltrating immune populations from middle-aged WT, *Tet2^+/-^* and *Dnmt3a^R^*^878^*^H/+^* CHIP animals. Frequency of ZsG+ mutant immune cells in the indicated population are depicted on the right y-axis. (G) Number of indicated immune cells in the tumor microenvironment normalized to the tumor weight /in grams, g). (H) Flow cytometry gating strategy for MHC-II expressing tumor-infiltrating macrophages (left panel) and fraction of MHC-II^high^ macrophages within those (right panel). For all panels, data is pooled from two independent experiments for *Tet2*-mutant and three experiments for *Dnmt3a*-mutant CHIP mice. Each dot represents one mouse. Statistical analysis was performed by using ordinary one-way ANOVA. Data is shown as mean with SEM. *p<0.05, **p<0.01, ***p<0.001, ****p<0.0001.

To assess age-related changes in tumor immunity, we compared young and middle-aged tumor-bearing mice across genotypes. In *Tet2^+/-^*-CHIP mice, aging increased circulating and TME myeloid cells (macrophages, neutrophils, and monocytes) and NK/NKT cells, while reducing CD4+, CD8+ T, and B cells. In contrast, WT and *Dnmt3a^R^*^878^*^H/+^*-CHIP mice showed fewer age-associated immune changes (**Figure S8A-D**), suggesting that aging affects more pronouncedly the immune responses of tumor-bearing *Tet2^+/-^*-CHIP mice. Of note, gating for ZsG+ cells revealed no significant expansion of mutant cells in any immune population compared with WT (**Figure 3F, right Y-axis**) or in PB following tumor inoculation (**Figure S5B-C**), indicating that tumor growth does not selectively promote CHIP expansion. Mutant cell frequencies were also not enriched in tumor leukocytes compared with PB, arguing against preferential recruitment of mutant cells to the TME (**Figure S5D**).

To further assess TME immunosuppression, we characterized macrophage polarization based on surface MHCII expression. *Tet2^+/-^*-CHIP tumors showed a significantly reduced frequency of MHCII-high “M1-like” macrophages compared with *Dnmt3aR878H/+*-CHIP or WT tumors (**Figure 3H, right panel**), which we also observed in middle-aged mice with a lower *Tet2*-mutant clonal fraction (**Figure S4R**). Collectively, consistent with the tumor bulk transcriptomic data, we observed a contrasting immune infiltrate between middle-aged mice with *Dnmt3a^R^*^878^*^H/+^*and *Tet2^+/-^*-CHIP, with the latter showing an overall increase in myeloid frequency, onset of immunosuppressive M2-like TAM polarization and reduced recruitment of adaptative immune cells.

### The tumor immune compartment in middle-aged *Tet2^+/-^*-CHIP mice exhibits an immunosuppressive phenotype with limited recruitment and effector function of cytotoxic CD8⁺ T cells

To uncover the transcriptomic changes in tumor-infiltrating immune cells from middle-aged WT, *Tet2^+/-^* and *Dnmt3a^-R^*^878^*^H/+^*-CHIP mice, we performed single-cell RNA sequencing on sorted ZsG+CD45+ mutant leukocytes from tumors of all three genotypes. Five major immune populations were identified: neutrophils, T/NK cells, B cells, dendritic cells, and monocytes-macrophages. Confirming the data from Figure 3, the monocyte-macrophage cluster represented the largest TME population (**Figure 4A, Table S2**). Next, we sub-clustered T cells and monocyte-macrophages clusters. We detected 9 main T cell sub-populations including CD8+ cytotoxic, CD8+ effector, naïve, CD4+ memory, CD4+ Tregs, CD8+ effector memory, CD8+ cytotoxic-proliferative, CD4+ proliferative Tregs and CD4+ resident memory T cells (**Figure 4B, upper panel, Table S3**). Cell number distribution by genotype clearly showed that in each T cell sub-population, *Tet2^+/-^*cells were underrepresented compared to WT and *Dnmt3a^R^*^878^*^H/+^* cells (**Figure 4B, lower panel**), in agreement with findings shown in Figure 3E-G. Similarly, sub-clustering of the monocyte-macrophage population revealed 8 distinct clusters; MHCII tumor-associated macrophages (MHCII-TAMs), angio-inflammatory TAMs, monocytic macrophages, lipid-associated TAMs (LA-TAMs), inflammatory TAMs (Inflam-TAMs), interferon-responsive TAMs (IFN-TAMs), monocytes and unconventional monocytes (**Figure 4C, upper panel, Table S4**). MHCII-TAMs and Angio-inflammatory TAMs constituted the majority of the myeloid compartments in TME, having distinct distributions across genotypes (**Figure 4C, lower panel**).

**Figure 4:**
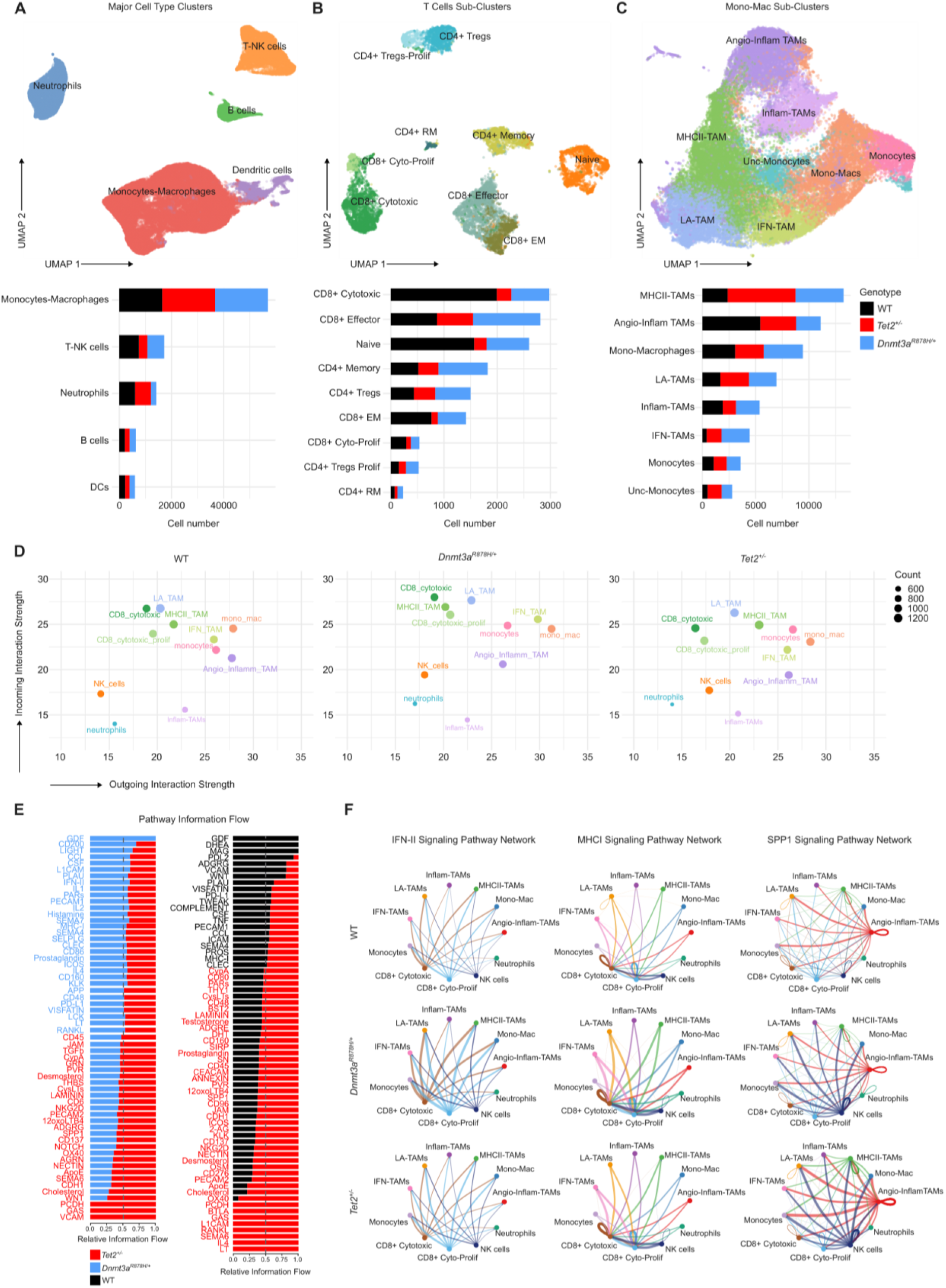
*Tet2*-mutant CHIP breast cancer-infiltrating immune cells show reduced cellular interactions, limited cytotoxic CD8+ T cell recruitment and effector status compared to WT and *Dnmt3a-*mutant CHIP. (A) Uniform manifold approximation and projection (UMAP) of major immune cell clusters from sorted CD45+ZsG+ immune cells from breast cancers of middle-aged WT, *Tet2^+/-^* and *Dnmt3a^R^*^878^*^H/+^* CHIP animals (upper panel) and distribution of genotypes and cell numbers in each cluster (lower panel). (B) UMAP of T cell sub-clustering (upper panel) and distribution of genotypes and cell numbers in each cluster (lower panel). (C) UMAP of monocytes/macrophage population (upper panel) and distribution of genotypes and cell numbers in each cluster (lower panel). (D) Cell type interaction strength from CellChat analysis showing incoming and outgoing interaction strength of indicated immune populations from WT, *Dnmt3a-* and *Tet2-*CHIP animals. (E) CellChat analysis showing the pathway information flow and the interaction strength between *Tet2*- and *Dnmt3a*-(left), and *Tet2*- and WT (right) chimeras. (F) Circle plots showing cell-cell interaction strength between major cell types from tumor microenvironment of animals for WT, *Dnmt3a ^fl-R^*^878^*^H/+^* and *Tet2^+/-^*CHIP animals for IFN-II (Type 2 interferon), MHC-I (major histocompatibility complex class I) and SPP1 (Secreted Phosphoprotein 1) signaling pathways.

We next characterized the cell-cell interaction between different immune clusters in tumors from WT, *Tet2^+/-^* and *Dnmt3a^fl-R^*^878^*^H/+^*-CHIP mice using CellChat. *Tet2^+/-^*-CHIP-bearing tumors showed reduced incoming interaction strength in CD8+ cytotoxic and proliferative T cells compared with WT and *Dnmt3aR878H/+*-CHIP, suggesting impaired responsiveness to TME cues, whereas MHCII-TAMs displayed increased outgoing interaction strength (**Figure 4D**). To gain deeper insights into the signaling pathways driving these changes, we identified altered pathways with distinct information flow between the conditions: *Dnmt3a^R^*^878^*^H/+^* vs *Tet2^+/-^*and WT vs *Tet2^+/-^*. We found decreased signaling via MHCI, IFNII and CCL pathways in tumors from *Tet2^+/-^*-CHIP compared to *Dnmt3a^R^*^878^*^H/+^*-CHIP, indicating impaired antigen presentation, immune activation, and leukocyte recruitment in tumors from *Tet2^+/-^*-CHIP mice. In line with this, *Tet2^+/-^*tumor-infiltrating immune cells showed increase in SPP1 pathway signaling compared to *Dnmt3a^R^*^878^*^H/+^* cells, consistent with an immunosuppressive TME (**Figure 4E, left panel**). Similarly, when comparing *Tet2^+/-^* to WT, MHCI signaling was increased in WT while SPP1 pathway interactions were enriched in *Tet2^+/-^*cells (**Figure 4E, right panel**). Circle plots further showed stronger MHCI signaling from *Dnmt3a^R^*^878^*^H/+^* macrophages and IFNII signaling from CD8+ cytotoxic T cells. Moreover, SPP1 signaling from angio-inflammatory and MHCII TAMs toward CD8+ T cells and other macrophages was enhanced in *Tet2^+/-^*-CHIP-bearing tumors (**Figure 4F**). Overall, *Tet2^+/-^*-CHIP-bearing tumors exhibited stronger immunosuppressive TAM signaling associated with impaired CD8+ T-cell responses, whereas *Dnmt3a^R^*^878^*^H/+^*-CHIP-bearing tumors showed enhanced immune-activating communication associated with increased cytotoxic T-cell activity.

### *Tet2^+/-^*MHCII-TAMs from middle-aged mice exhibit immunosuppressive phenotype and increased expression of *Cxcl2* and *Il1a/b*

To identify targetable drivers of the immunosuppressive polarization of TAMs, we sub-clustered MHCII-TAMs, revealing three distinct clusters. Cluster 1 had the lowest *Tet2^+/-^* and highest *Dnmt3a^R^*^878^*^H/+^*contribution while cluster 3 showed the opposite pattern (**Figure 5A, B**). Notably, top markers of cluster 3 included immunosuppressive TAM-associated genes such as *Cxcl2, Clec4e, Slc7a11, Cd9* and *Spp1*^23–27^ (**Figure 5C**). Compared with cluster 1, cluster 3 showed reduced Hallmark IFNα and IFNγ responses and increased NF-κB signaling (**Figure 5D, E**), which was supported by decreased expression of IFNγ-inducible T cell chemoattractant encoding genes *Cxcl9/10* (**Figure 5F**). Consistent with our macrophage polarization data, a myeloid-derived suppressor cell (MDSC) gene signature^28^ was enriched in cluster 3 (**Figure 5G**), confirming its immunosuppressive transcriptional profile. Intersection of the MDSC and Hallmark NF-κB signatures identified 10 shared genes, including *Il1b* and *Cxcl2*, both implicated in M2-like immunosuppressive macrophage polarization (**Figure 5H**) ^23,29^. UMAPs depicting the expression of *Cxcl2*, *Il1a* and *Il1b* confirmed higher expression of these genes in cluster 3 (**Figure 5I**). Similar immunosuppressive signatures, including increased *Cxcl2* and *Il1a/b*, were observed in the second-largest TAM cluster, TAMs Angio-inflammatory (**Figure S9A-H**). In sum, *Tet2^+/-^*-enriched MHCII and Angio-inflammatory TAMs displayed an immunosuppressive program characterized by increased *Il1a/b* and its downstream target *Cxcl2*, highlighting these pathways as potential therapeutic vulnerabilities.

**Figure 5:**
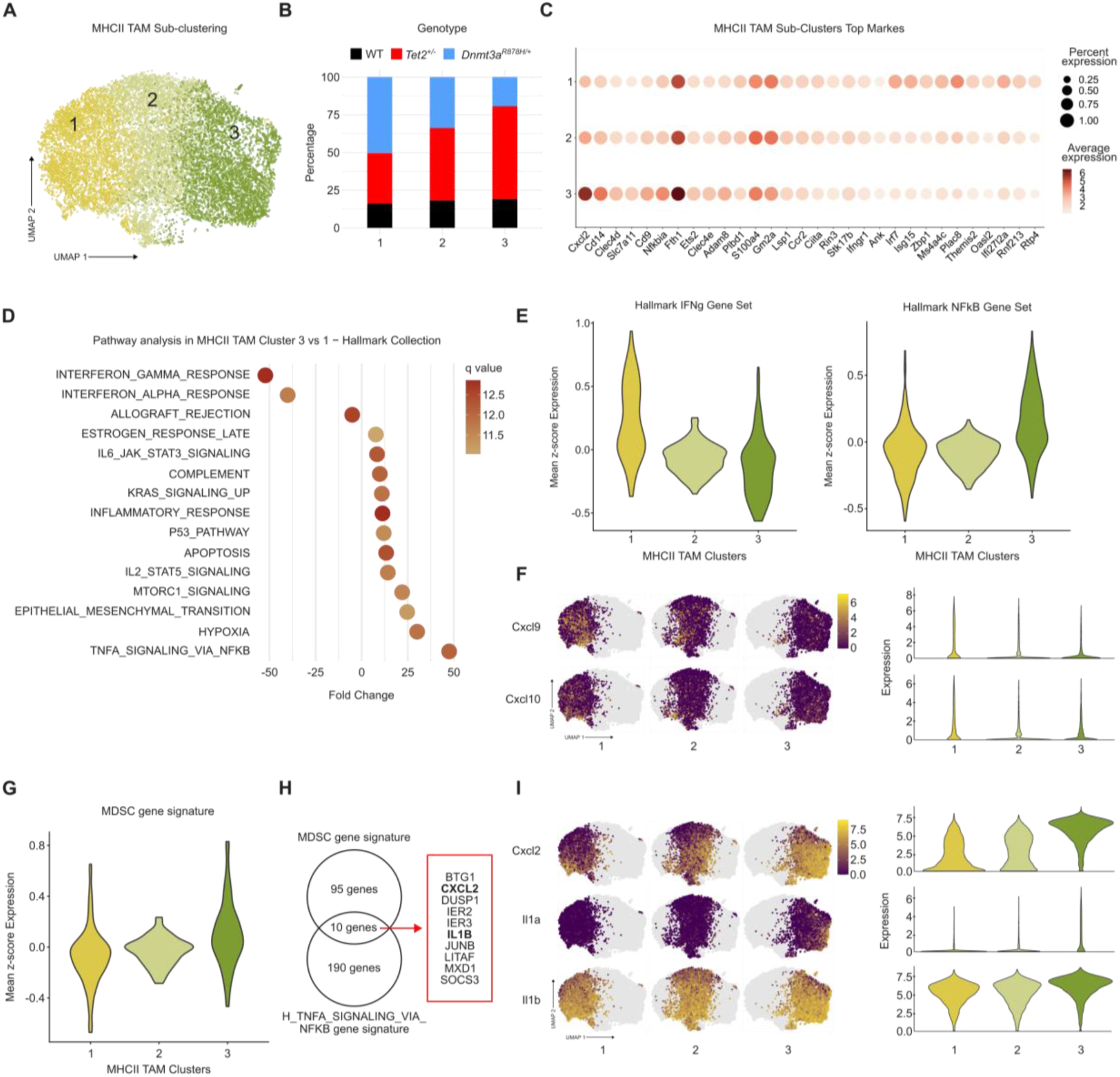
The *Tet2*-mutant MHCII-TAM enriched clusters show increased NF-kB activation, immunosuppressive signatures and higher IL-1 / Cxcl2 levels compared to WT and *Dnmt3a*-mutant TAMs. (A) UMAP showing the sub-clustering of MHC-II TAM populations. (B) Distribution of genotypes in each sub-cluster. (C) Dot plot showing the expression of top markers in each sub-cluster of MHCII-TAMs. (D) Pathway analysis in MHCII-TAM sub-clusters 3 vs 1 in the Hallmark dataset. (E) Violin plots showing the mean z-score for expression of genes in the Hallmark IFNγ (left) and Hallmark NF-kB (right) gene sets in MHCII-TAM sub-clusters. (F) UMAP of sub-clusters of MHC-II TAM population showing the expression of Cxcl9 (upper panel), Cxcl10 (lower panel) and respective quantification in violin plots. (G) Violin plots showing the mean z-score for expression of genes in the myeloid-derived suppressor (MDSC) signature in the MHC-II TAM populations. (H) Venn diagram depicting the intersection of genes from the MDSC set and the Hallmark-NFkB set. The common 10 genes are depicted. (I) UMAP of sub-clusters of MHC-II TAM population showing the expression of Cxcl2 (upper panel), IL-1α (middle panel) and IL-1β (lower panel) and respective quantification in violin plots.

### CXCR2 and IL-1R1 pharmacologic blockage in middle-aged *Tet2^+/-^*-CHIP mice reduces breast tumor progression via attenuation of immunosuppressive macrophage polarization

To validate the IL-1 pathway as a potential driver of a pro-tumoral immune microenvironment, we first evaluated the effect of systemic IL-1α and -β administration on tumor growth in young WT mice using the orthotopic E0771 model (**Figure S10A-B**). IL-1α, but not IL-1β, promoted breast tumor growth and increased lung metastatic burden compared with PBS-treated controls (**Figure S10C-H**). Moreover, IL-1α expanded myeloid populations in PB and BM (**Figure S10I-J**) but had a minor impact on tumor immune composition (**Figure S10K-M**). The tumor-promoting effect of IL-1α was unlikely to result from direct sensing of IL-1α by E0771 cells, as in vitro exposure to low and high doses of IL-1α and -β resulted in either no effect or reduced tumor cell numbers, respectively (**Figure S10N**). These findings identify IL-1α as a driver of E0771 tumor progression in vivo, likely through modulation of TME immune functions, consistent with previous findings^30^. Next, we investigated whether breast cancer progression in middle-aged *Tet2^+/-^*-CHIP mice could be therapeutically modulated by targeting IL-1R1 and CXCR2, the cognate receptors for IL-1 and CXCL2, respectively. Following induction of high *Tet2*^+/-^chimerism and 10 months of aging, E0771 cells were orthotopically transplanted, and mice were treated one week later with the clinically approved IL-1R1 antagonist Anakinra or the CXCR2 antagonist SB225002 (**Figure 6A-B**). Longitudinal measurement of tumor growth revealed that both antagonists reduced breast tumor progression in middle-aged *Tet2^+/-^*-CHIP mice to the level of WT controls, demonstrating that the tumor promoting effect of *Tet2^+/-^*-CHIP in middle-aged mice can be abrogated by blocking the CXCL2 or IL-1 pathways (**Figure 6C-F**). Both interventions also reduced lung metastatic burden, although this effect did not reach statistical significance (**Figure S11A**).

**Figure 6:**
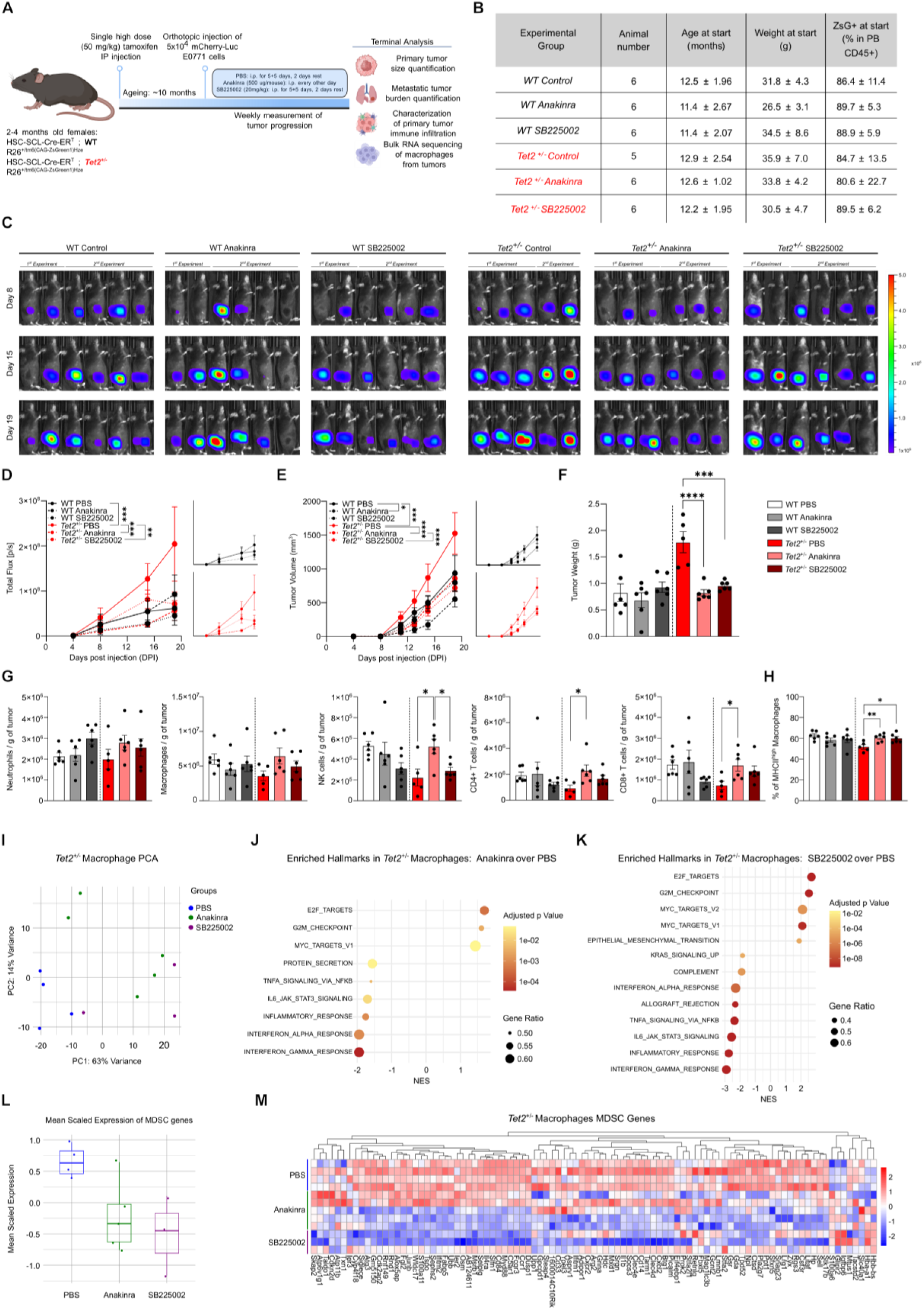
IL-1R1 and CXCR2-pathway inhibition reduces breast cancer growth to WT background level in middle-aged mice with *Tet2*-mutant CHIP. (A) Graph depicting experimental outline. (B) Graph depicting animal numbers, age, weight and peripheral blood chimerism (CD45+ ZsG+) of each experimental group. (C) *In vivo* growth of mCherry-Luciferase expressing E0771 breast cancer over time by IVIS imaging of WT and *Tet2^+/-^*CHIP animals treated with PBS, Anakinra or SB225002. (D) Tumor growth curves (left panel) and individual curves (right panel) by quantification of luminescent intensity from images in (C). (E) Tumor growth curves (left panel) and individual curves (right panel) by quantification tumor volume in mm^3^(right). (F) Tumor weight measurement in grams at termination of experiment at day 20. (G) Flow cytometry analysis showing the number of indicated immune cells in the tumor microenvironment per gram of tumor. (H) Fraction of MHC-II^high^ in total macrophages. (I) Principal component analysis (PCA) of bulk RNA sequencing of macrophages from tumor microenvironment of *Tet2^+/-^*CHIP animals with PBS, Anakinra or SB225002 treatment. (J) Gene set enrichment analysis (GSEA) of macrophages showing enriched hallmark datasets from *Tet2^+/-^*CHIP animals, treated with Anakinra vs PBS. (K) GSEA of macrophages showing enriched hallmark datasets from *Tet2^+/-^*CHIP animals, treated with SB225002 vs PBS. (L) Box plots showing the mean scaled expression of genes in the myeloid-derived suppressor (MDSC) signature in PBS, Anakinra and SB225002 treated *Tet2^+/-^*CHIP animals. (M) Heatmap showing the expression of genes for MDSC signature from *Tet2^+/-^*macrophages. For panels (C-H), data is pooled from two independent experiments. Statistical analysis was performed by using two-way ANOVA (D, E) and ordinary one-way ANOVA within each genotype (F, G, H). Data is shown as mean with SEM. *p<0.05, **p<0.01, ***p<0.001, ****p<0.0001.

Focusing on treatment-induced immune alterations, Anakinra prevented tumor-induced myeloid expansion in PB and reduced it in BM of *Tet2^+/-^*-CHIP mice, confirming IL-1 as a driver of cancer-adapted myelopoiesis in this model (**Figure S11B, C**). Of note, neither short-term treatment affected the *Tet2*-mutant clonal size measured in PB or BM leukocytes (**Figure S10D-G**), unlike the reduction previously observed in tumor-free aged mice treated with Anakinra for 2 months^21^. In the TME, Anakinra administration increased NK, CD4+ T, CD8+ T and DN T cell infiltration, while SB225002 administration led to a non-significant increase in CD8+ T cells, suggesting a partial reversion of the immunosuppressive TME quality (**Figure 6G, Figure S11H-J**). In agreement, both treatments increased MHCII expression in TAMs, suggesting a shift towards an anti-tumorigenic M1-like phenotype (**Figure 6H**). To define the transcriptional programs underlying this shift, we performed bulk RNA-sequencing of WT and *Tet2^+/-^*TAMs exposed to Anakinra and SB225002. Focusing our analysis on *Tet2^+/-^*macrophages, PCA revealed a separation between macrophages isolated from PBS-treated mice and those from Anakinra/SB225002-treated mice (**Figure 6I, Table S5**). GSEA revealed that *Tet2^+/-^*macrophages exposed to either treatment exhibited reduced expression of interferon response, JAK-STAT3 and NF-kB pathway genes, while displaying increased enrichment of MYC and E2F target gene programs, suggesting diminished inflammatory signaling accompanied by enhanced proliferative activity (**Figure 6J, K**). Finally, to evaluate the impact of treatments on the immunosuppressive properties of tumor-infiltrating *Tet2^+/-^*macrophages, we evaluated the expression of the previously used MDSC signature (**Figure 5F**). Strikingly, we observed that both Anakinra and SB225002 treatments significantly reduced the mean expression of MDSC genes, with a clear decrease in expression of well-established macrophage associated immunosuppressing genes like *Pla2g7, Ppt1, Cd84, Csf3r, Il4ra, Slc40a1, Cxcr2, Il1b and Cxcl2*^23,29,31–36^ (**Figure 6L, M**). Overall, these findings indicate that IL-1 and CXCL2 over-produced by *Tet2^+/-^*TAMs contribute to breast cancer progression by promoting an immunosuppressive M2-like phenotype, thereby limiting anti-tumor immunity and cytotoxic T-cell responses.

## DISCUSSION

Recent large patient cohort studies have linked CHIP, detected in blood and tumors, to increased cancer incidence and mortality^6,8,37^. In breast cancer, CHIP is associated with increased incidence and chemotherapy-related toxicities, although its effects on disease progression remain unclear^15–20,37^. Here, using inducible mouse models of *Dnmt3a*- and *Tet2*-mutant CHIP combined with syngeneic orthotopic breast cancer models, we demonstrate that the impact of CHIP on tumor progression is determined by the mutant gene, clonal size, and host age. Indeed, *Dnmt3a*-mutant CHIP delayed breast cancer progression independently of age, whereas *Tet2*-mutant CHIP promoted progression specifically in middle-aged mice, with effects proportional to clonal size. These opposing effects reflected distinct remodeling of TME: *Dnmt3a*-mutant CHIP generated an interferon-rich TME characterized by increased MHC expression in tumor cells and TAMs and enhanced T-cell recruitment and activation, whereas *Tet2*-mutant CHIP promoted an immunosuppressive TAM phenotype with reduced T-cell recruitment and activation. Importantly, targeting inflammatory factors overproduced by aged *Tet2*-mutant TAMs, including IL-1 and CXCL2, reduced TAM immunosuppression and tumor growth to levels observed in WT mice.

A key finding of this study is that *Tet2*-mutant CHIP accelerates breast tumor progression and metastasis with aging, whereas *Dnmt3a*-mutant CHIP protects against primary tumor growth, demonstrating that CHIP genotype determines disease outcome. This aligns partially with a recent study of 125 breast cancer patients treated with radiotherapy, in which high-burden (VAF >10%) non-*DNMT3A* CHIP was associated with shorter distant metastasis-free survival, whereas *DNMT3A*-mutant CHIP showed outcomes comparable to no-CHIP controls^19^. Consistent with these findings, *TET2*-mutant CHIP has also been reported to be more frequently associated with tumor-infiltrating CH than *DNMT3A*-mutant CHIP, suggesting a greater impact of *TET2* mutations on the TME^8^. While the mechanisms underlying this difference were not clarified in these prior studies, our findings indicate that *Tet2*-mutant CHIP profoundly alters TME immune competence. *Tet2*-mutant TAMs exhibit increased production of IL-1α/β and CXCL2, associated with NF-κB activation, enhanced myeloid recruitment and immunosuppressive TAM polarization, reduced MHCII expression, and impaired CD4⁺ and CD8⁺ T-cell recruitment and activation. These findings align with previous reports of increased IL-1 and CXCL2 production by *Tet2*-mutant macrophages and with evidence that loss of TET2 promotes immunosuppressive macrophage polarization through altered regulation of NF-κB target genes^21,38–40^. Moreover, clinical and preclinical studies further support an association between *Tet2* loss, increased tumor-infiltrating myeloid cells, an immunosuppressed TME, angiogenesis, and tumor growth^8,9,11,41^. Contrasting our findings in a therapeutic setting, studies using bone marrow transplantation-based *Tet2*-mutant CHIP models reported enhanced TAM antigen presentation and improved responses to immune checkpoint blockade^42,43^. These differences likely reflect the distinct tumor and CHIP models used and emphasize the role of tumor-specific microenvironmental cues in shaping CHIP-associated immune responses. Collectively, our data establishes that, in syngeneic orthotopic breast cancer models, *Tet2*-mutant CHIP promotes tumor progression through IL-1/CXCL2-driven immunosuppressive TAM polarization.

Opposing to *Tet2*-mutant TAMs, *Dnmt3a*-mutant TAMs maintained an interferon-responsive transcriptome, increased MHCII expression, and higher production of the T-cell chemoattractants CXCL9 and CXCL10. This was associated with enhanced recruitment and activation of IFNγ-producing CD8⁺ and CD4⁺ T cells and reduced tumor progression. Consistent with a protective role, *DNMT3A*-mutant CHIP has been associated with improved survival in metastatic colorectal cancer, with clonal size correlating with overall survival in the FIRE-3 trial^44^. Subsequent studies suggested that this effect may be mutation-specific, with the R882 hotspot (equivalent to the R878H mutation used here) being protective, whereas non-R882 mutations may be pro-tumoral^45^. Although we could not definitively establish the mechanism underlying the protective immune response, *Dnmt3a*-mutant CHIP was associated with an IFNγ-responsive TME, contrasting with the NF-κB-dominated TME of *Tet2*-mutant CHIP. DNMT3A has been implicated in regulation of *IFNG* expression, and *DNMT3A*-deficient CD4⁺ T cells exhibit increased *IFNG* expression following activation^46^. Consistent with this, donor-derived *DNMT3A* CHIP in hematopoietic stem cell (HSC) transplantation is associated with increased graft-versus-host disease and reduced leukemia relapse^47–49^. Thus, increased IFNγ production by tumor-infiltrating *Dnmt3a*-mutant T cells could contribute to the protective phenotype, although this remains unproven. However, whereas *DNMT3A* mutations in T cells may arise from mutant HSC-driven T cell reconstitution after transplantation, they are rare in non-transplant settings, particularly in the TME^8^, and unlikely to substantially influence clinical outcomes. Thus, as in most CHIP models, *Dnmt3a* loss across immune cells, including T cells, may introduce effects that do not fully reflect the clinical setting. Nevertheless, the identified mutation-specific impacts in the TME might underly the currently contradictory data concerning the impact of CHIP on breast cancer patient outcomes, particularly when *DNMT3A*- and *TET2*-mutant CHIP are considered as a single entity. Larger breast cancer patient cohorts will be required to define these mutation-specific effects clinically^17^.

Another relevant finding is that the tumor-promoting effect of *Tet2*-mutant CHIP occurs specifically in middle-aged mice, indicating that its impact depends not only on the mutation itself but also on age-dependent systemic changes. Aging is accompanied by immune dysfunction, chronic inflammation, myeloid skewing, impaired macrophage function, and increased T- and B-cell exhaustion and clonal restriction^50^. These age-associated changes create a cellular and molecular environment that both shapes immune responses and influences mutant HSC selection. Indeed, microbiota-driven IL-1 signaling represents a key age-associated selective pressure for *Tet2*-mutant HSCs^21,51^, whereas infection-induced IFNγ can promote selection of *Dnmt3a*-mutant HSCs^52^.

Thus, CHIP arises mainly in aging hematopoietic systems and can further exacerbate hematopoietic aging by amplifying inflammatory signaling and dysregulated cytokine production. In our model, aging modestly increased breast cancer progression in WT mice, consistent with previous observations^53^, but had a markedly greater effect in *Tet2*-mutant CHIP mice, where tumor growth was accompanied by exacerbated myelopoiesis and increased TAM and TAN accumulation. We propose that this enhanced phenotype results from increased IL-1 and downstream CXCL2 production by aged *Tet2*-mutant TAMs, which may be less pronounced in young mice. One potential driver is the increase in microbiota-derived systemic signals with age^51^, also observed following inflammation-induced intestinal permeability in mice lacking hematopoietic *Tet2*^54^. Thus, aging and *Tet2*-mutant CHIP may synergistically increase exposure to microbiota-derived signals, enhancing IL-1 production by aged mutant TAMs. Alternatively, aging may promote progressive seeding of *Tet2*-mutant monocyte-derived macrophages into the breast fat pad before tumor development, generating a pre-existing immunosuppressive environment with a higher burden of mutant cells.

Finally, targeting IL-1 signaling in aged tumor-bearing mice with *Tet2*-mutant CHIP significantly reduced primary tumor growth, decreased TAM immunosuppressive polarization, and restored tumor-infiltrating CD4⁺, CD8⁺ T, and NK cells to WT levels. These findings are consistent with evidence implicating IL-1 signaling in breast cancer immunosuppression^55,56^ and identify increased IL-1 production by *Tet2*-mutant TAMs as a targetable driver of age-associated tumor promotion. This is further supported by clinical data linking tumor-infiltrating CH to increased *IL1A* expression and an IL-1A transcriptional program in tumors from older patients^9^. Whether IL-1 targeting remains effective during cancer treatment is unknown, although chemotherapy may further enhance IL-1α production and immunosuppression^30^. CXCR2 inhibition also reduced tumor growth in aged *Tet2*-mutant CHIP mice, although its effects on the TME were more moderate. Given the established role of the CXCL2-CXCR2 axis in myeloid recruitment and its contribution to breast cancer progression^57,58^, these findings suggest a complementary role for this pathway. Indeed, both IL-1R1 and CXCR2 inhibition reduced inflammatory/NF-κB signatures and immunosuppressive MHCII low TAM polarization, while *Cxcr2*, *Cxcl2*, and *Il1b* were downregulated following either treatment. This suggests that IL-1R1- and CXCR2-signaling pathways create inflammatory loops in the *Tet2*-mutant macrophage population, which drives these cells towards immunological dysfunction and facilitates tumor progression in *Tet2*-mutant CHIP carriers. Whether targeting either IL-1R1, CXCR2 or both in combination with standard treatment will improve breast cancer patient outcomes, particularly in TET2-mutant CHIP cases, remains to be determined.

## RESOURCE AVAILABILITY

The datasets generated from the current study are available from the corresponding author on reasonable request. The bulk and single-cell RNA sequencing data sets generated in this study will be deposited in Gene Expression Omnibus (GEO) and will be made publicly available upon publication of the manuscript.

## ACKNOWLEDGEMENTS

The authors thank Syndi Uhlig and Angelina Oestmann for technical assistance with laboratory and mouse colony management. The authors also thank Lennart Opitz and the Functional Genomic Centre Zurich team as well as Susanne Dettwiler and Fabiola Prutek from the Department of Pathology and Molecular Pathology for technical assistance.

This work was supported by the Swiss National Science Foundation (320030_219676/1) (M.G.M.).

## AUTHOR CONTRIBUTIONS

N.G.G designed and performed experiments, analyzed data, and wrote the manuscript; K.Z., J.F. and F.P. performed experiments and analyzed data; S.B. supervised research; F.C. performed experiments, analyzed data, designed and supervised research and wrote the manuscript. M.G.M. designed and supervised research and wrote the manuscript.

## DECLARATION REGARDING THE USE OF AI

During the preparation of this work, the authors used ChatGTP to reduce manuscript word count. After using this tool, the authors reviewed and edited the content as needed and take full responsibility for the content of the publication.

## FIGURES

**Supplementary Figure 1:**
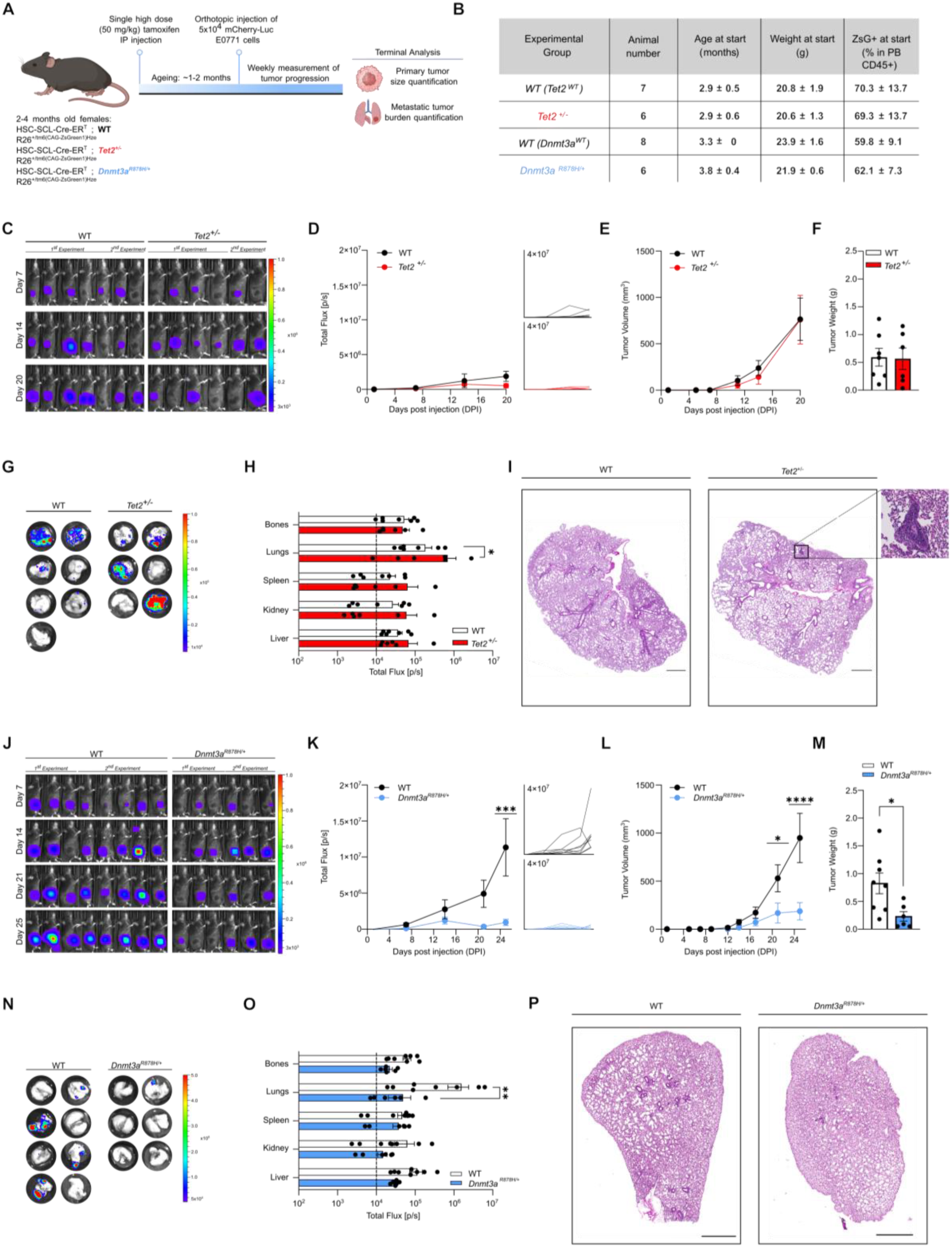
*Dnmt3a*-mutant CHIP reduces breast cancer growth in young mice, while *Tet2*-mutant CHIP has no significant impact on cancer growth. (A) Graph depicting experimental outline. (B) Graph depicting animal numbers, age, weight and peripheral blood chimerism (CD45+ ZsG+) of each experimental group. (C) *In vivo* growth of orthotopic injected, mCherry-Luciferase expressing E0771 breast tumor cell line over time, evaluated by IVIS imaging of WT and *Tet2^+/-^*CHIP animals. (D) Tumor growth curves (left panel) and individual curves (right panel) determined by quantification of luminescent intensity from images in (C). (E) Tumor volume measurements over time in mm^3^. (F) Tumor weight measurement in grams at terminal analysis. (G) *Ex vivo* IVIS images of lungs showing level of metastasis in WT and *Tet2^+/-^*CHIP animals. (H) Ex vivo quantification of luminescence intensity in indicated organs of WT and *Tet2^+/-^*CHIP animals (G). Dashed line represents the average background signal. (I) Representative HE stained lung section, scale bar 800µm, higher magnification of the area of interest. (J) *In vivo* growth of orthotopic injected, mCherry-Luciferase expressing E0771 breast tumor over time by IVIS imaging of WT and *Dnmt3a ^fl-R^*^878^*^H/+^*CHIP animals. (K) Tumor growth curves (left panel) and individual curves (right panel) determined by quantification of luminescent intensity from images in (J). (L) Tumor volume measurements over time in mm^3^ from WT and *Dnmt3a^R^*^878^*^H/+^* CHIP animals. (M) Tumor weight measurement in grams at terminal analysis. (N) *Ex vivo* IVIS images of lungs showing level of metastasis in WT and *Dnmt3a^R^*^878^*^H/+^* CHIP animals. (O) *Ex vivo* quantification of luminescent intensity in indicated organs from WT and *Dnmt3a ^fl-R^*^878^*^H/+^*CHIP animals. Dashed line represents the average background signal (P) Representative HE staining of lung sections, scale bar 800µm. For all panels, data is pooled from two independent experiments. Statistical analysis was performed by using two-way ANOVA (D, E, H, K, L, O) and unpaired t test (F, M). Data is shown as mean with SEM. *p<0.05, **p<0.01, ***p<0.001, ****p<0.0001

**Supplementary Figure 2:**
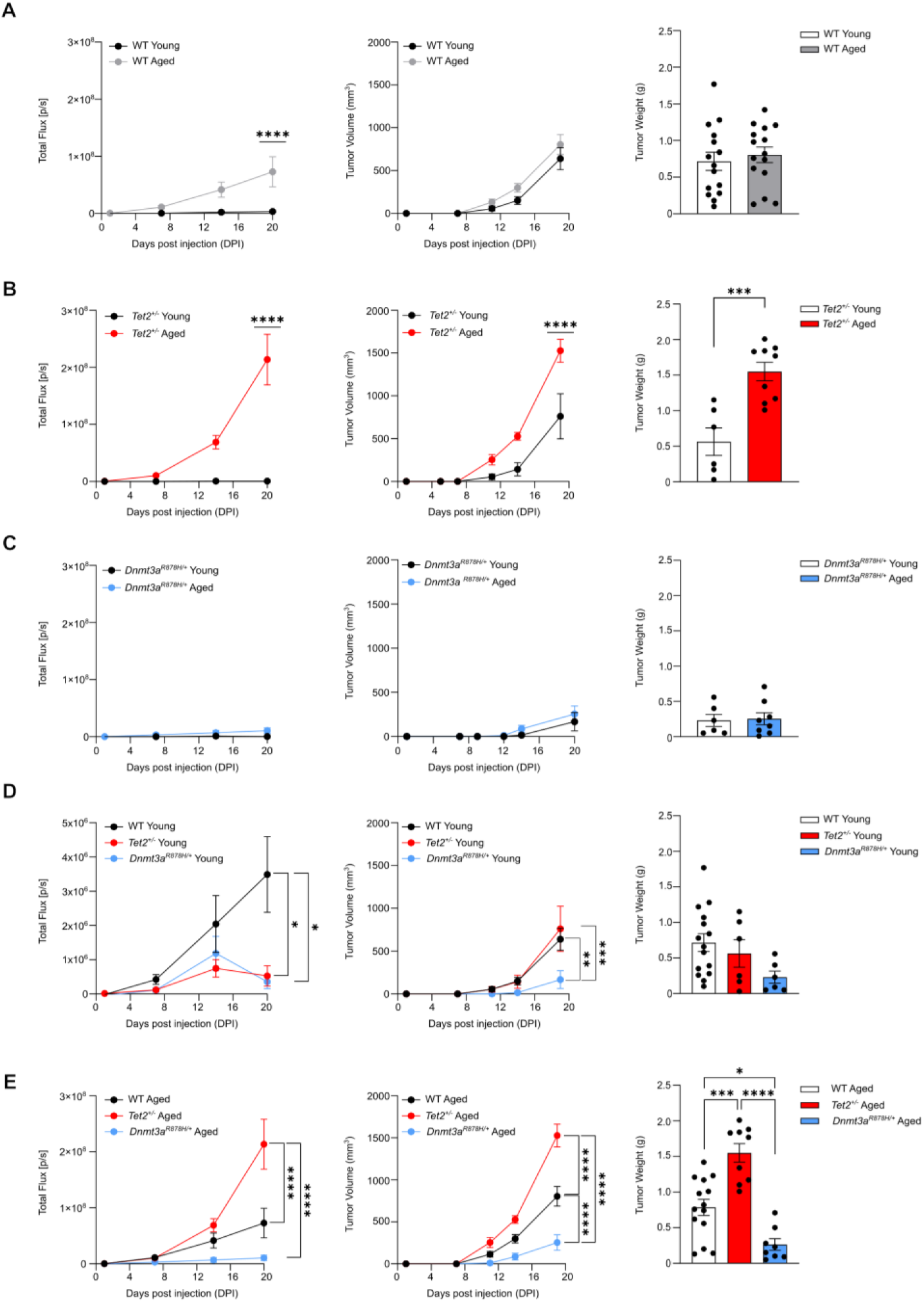
Impact of aging on breast cancer progression within each genotype. (A) Tumor growth curves of orthotopic injected, mCherry-Luciferase expressing E0771 breast cancer cell line over time, determined by quantification of luminescent intensity by IVIS imaging of WT young and middle-aged animals (left panel). Tumor volume measurements over time in mm^3^ (middle panel). Tumor weight measurement in grams at terminal analysis (right panel). (B) Tumor growth curves of orthotopic injected, mCherry-Luciferase expressing E0771 breast tumor cell line over time determined by quantification of luminescent intensity by IVIS imaging of *Tet2^+/-^*CHIP young and middle-aged animals (left panel). Tumor volume measurements over time in mm^3^ (middle panel). Tumor weight measurement in grams (right panel). (C) Tumor growth curves of orthotopic injected, mCherry-Luciferase expressing E0771 breast tumor cell line over time determined by quantification of luminescent intensity by IVIS imaging of *Dnmt3a ^fl-R^*^878^*^H/+^* CHIP young and middle-aged animals (left panel). Tumor volume measurements over time in mm^3^ (middle panel). Tumor weight measurement in grams (right panel). (D) Tumor growth curves of orthotopic injected, mCherry-Luciferase expressing E0771 breast cancer cell line over time determined by quantification of luminescent intensity by IVIS imaging young WT, *Tet2^+/-^*and *Dnmt3a^R^*^878^*^H/+^* CHIP animals (left panel). Tumor volume measurements over time in mm^3^ (middle panel). Tumor weight measurement in grams (right panel). (E) Tumor growth curves of orthotopic injected, mCherry-Luciferase expressing E0771 breast cancer cell line over time determined by quantification of luminescent intensity by IVIS imaging middle-aged WT, *Tet2^+/-^* and *Dnmt3a^R^*^878^*^H/+^* CHIP animals (left panel). Tumor volume measurements over time in mm^3^ (middle panel). Tumor weight measurement in grams (right panel). For all panels, data is pooled from Fig. 1D, E, F, L M, N and Supp. Figure 1D, E, F, K, L, M. Statistical analysis was performed by using two-way ANOVA for growth curves, unpaired t test for tumor weights in panel A, B, C and ordinary one-way ANOVA for tumor weights in panel D, E. Data is shown as mean with SEM. *p<0.05, **p<0.01, ***p<0.001, ****p<0.0001.

**Supplementary Figure 3:**
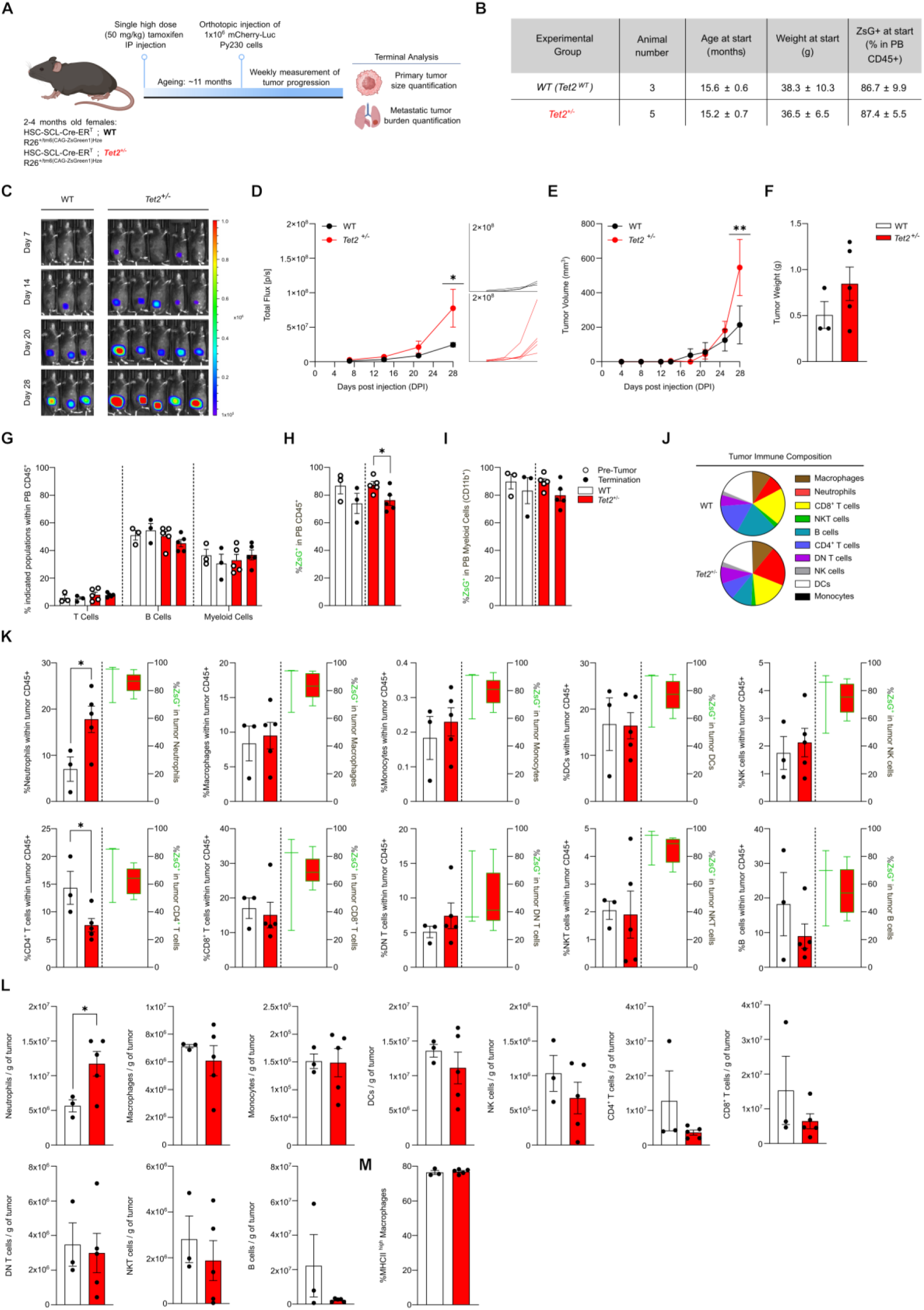
*Tet2*-mutant CHIP promotes Py230 tumor progression in middle-aged mice. (A) Graph depicting experimental outline. (B) Graph depicting animal numbers, age, weight and peripheral blood chimerism (CD45+ ZsG+) of each experimental group. (C) *In vivo* growth of orthotopic injected, mCherry-Luciferase expressing Py230 breast tumor cell line over time, evaluated by IVIS imaging of WT and *Tet2^+/-^*CHIP animals. (D) Tumor growth curves (left panel) and individual curves (right panel) determined by quantification of luminescent intensity from images in (C). (E) Tumor volume measurements over time in mm^3^. (F) Tumor weight measurement in grams at terminal analysis. (G) Frequency of T cells, B cells and myeloid cells (CD11b+) in peripheral blood (PB) within CD45+ leukocytes before orthotopic breast cancer injection and at terminal analysis. (H) Frequency of ZsG+ mutant immune cells in PB within CD45+ leukocytes before tumor injection and at the terminal analysis from WT and *Tet2^+/-^*CHIP animals. (I) Frequency of ZsG+ mutant immune cells in PB within CD45+CD11b+ myeloid population before breast cancer cell injection and at terminal analysis. (J) Tumor immune composition as a percentage of indicated immune populations in CD45+ leukocytes in tumors. (K) Flow cytometric analysis of indicated tumor infiltrating immune populations from middle-aged WT and *Tet2^+/-^*CHIP mice. Frequency of ZsG+ mutant immune cells in the indicated population are depicted on the right y-axis. (L) Number of immune cells in the tumor microenvironment normalized to the gram of tumor for indicated populations. (M) Fraction of MHC-II^high^ in total tumor macrophages from WT and *Tet2^+/-^*CHIP animals. Statistical analysis was performed by using two-way ANOVA (D, E, G) and unpaired t test (F, H, I, K-M). Data is shown as mean with SEM. *p<0.05, **p<0.01, ***p<0.001, ****p<0.0001.

**Supplementary Figure 4:**
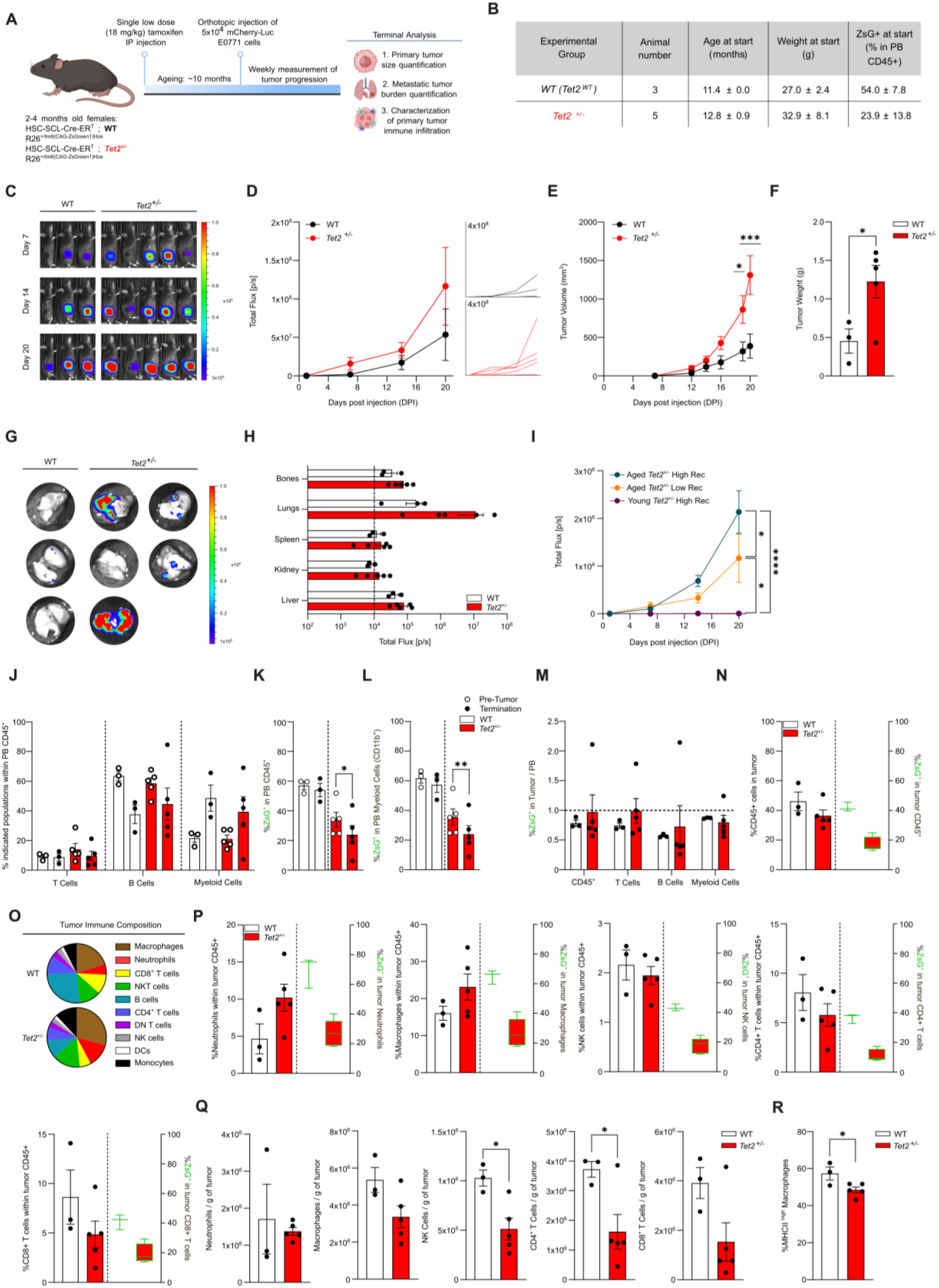
*Tet2*-mutant CHIP with a lower clonal hematopoietic fraction is sufficient to enhance breast cancer growth in middle-aged mice. (A) Graph depicting experimental outline. (B) Graph depicting animal numbers, age, peripheral blood chimerism (CD45+ ZsG+) of each experimental group. (C) *In vivo* growth of orthotopic injected, mCherry-Luciferase expressing E0771 breast tumor over time, evaluated by IVIS imaging of WT and *Tet2^+/-^*CHIP animals. (D) Tumor growth curves (left panel) and individual curves (right panel) by quantification of luminescent intensity from images in (C). (E) Tumor volume measurements over time in mm^3^. (F) Tumor weight measurement in grams at terminal analysis. (G) *Ex vivo* IVIS images of lungs showing level of metastasis in WT and *Tet2^+/-^*CHIP animals. (H) Quantification of luminescent intensity of *ex vivo* images by IVIS showing level of metastasis in distal organs from WT and *Tet2^+/-^*CHIP animals. Dashed line represents the average background signal. (I) Tumor growth curves by quantification of luminescent intensity from young *Tet2^+/-^*CHIP animals with high recombination, middle-aged *Tet2^+/-^*animals with low recombination and middle-aged *Tet2^+/-^*CHIP animals with high recombination. Data is pooled from the experiments shown in Supp. Fig. 1D, Supp. Fig 3D and Fig 1D. (J) Frequency of T cells, B cells and myeloid cells in peripheral blood (PB) within CD45+ leukocytes before tumor injection and at the terminal analysis. (K) Frequency of ZsG+ mutant immune cells in PB within CD45+ leukocytes before tumor injection and at the terminal analysis from WT and *Tet2^+/-^*CHIP animals. (L) Frequency of ZsG+ mutant immune cells in PB within CD45+CD11b+ myeloid population before breast cancer cell injection and at the terminal analysis. (M) Relative enrichment of ZsG+ mutant immune cell frequency in total CD45+ leukocytes, T cells, B cells and myeloid cells in tumors compared to PB from WT and *Tet2^+/-^*CHIP animals. (N) Percentage of total CD45^+^ leukocytes in total live cells of tumor microenvironment in WT and *Tet2^+/-^*CHIP animals by flow cytometric analysis. Frequency of ZsG+ mutant immune cells in the indicated population are depicted on the right y-axis. (O) Tumor immune composition as a percentage of indicated immune populations in CD45+ leukocytes in tumor. (P) Flow cytometric analysis of indicated tumor infiltrating immune populations from middle-aged WT and *Tet2^+/-^-*CHIP mice. Frequency of ZsG+ mutant immune cells in the indicated population are depicted on the right y-axis. (Q) Number of immune cells in the tumor microenvironment normalized to the gram of tumor for indicated populations. (R) Fraction of MHC-II^high^ in total tumor macrophages from WT and *Tet2^+/-^*animals. Statistical analysis was performed by using two-way ANOVA (D, E, H, I, J, M), paired t test (K, L) and unpaired t test (F, N, P, Q, R). Data is shown as mean with SEM. *p<0.05, **p<0.01, ***p<0.001, ****p<0.0001.

**Supplementary Figure 5:**
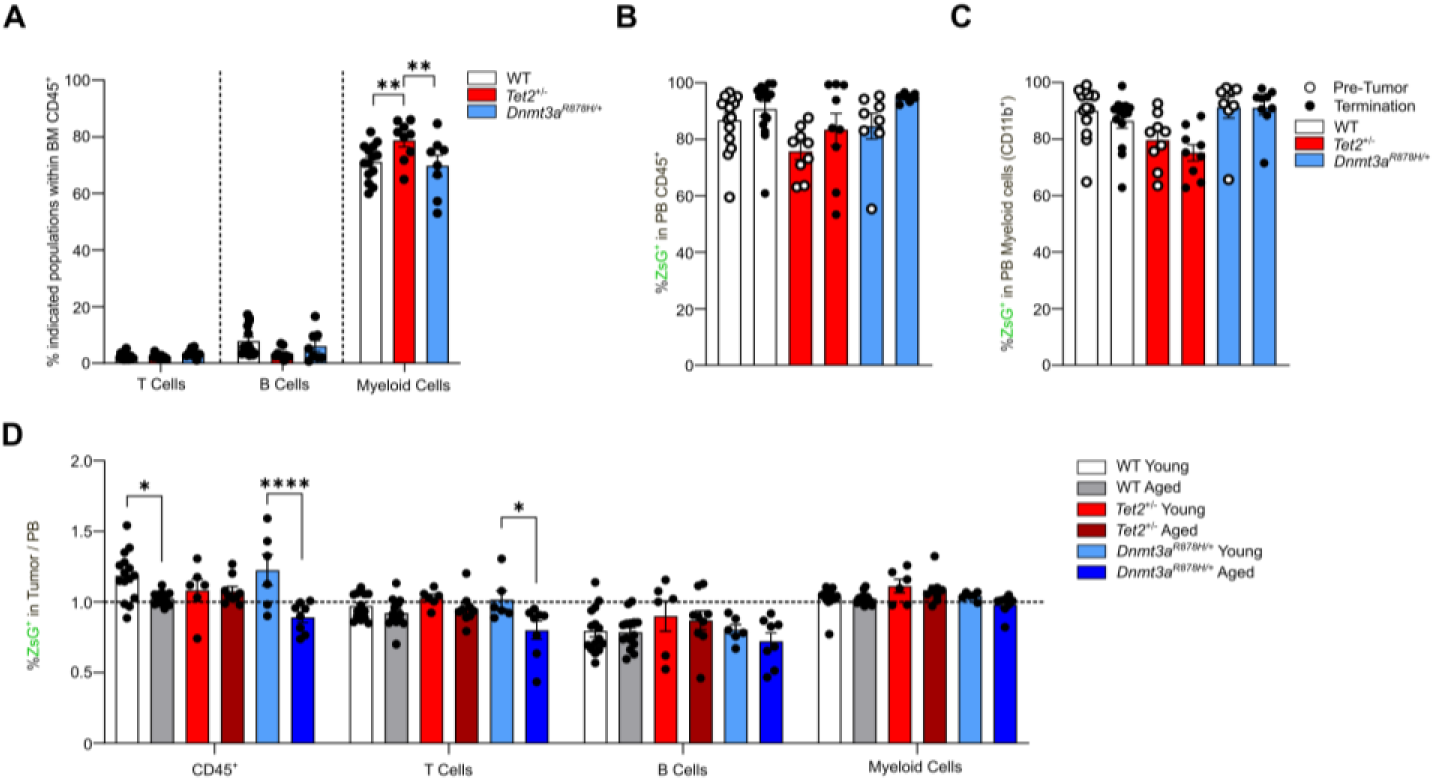
Immune characterization of tumors from middle-aged mice with WT, *Tet2*-or *Dnmt3a*-mutant CHIP. (A) Frequency of T cells, B cells and myeloid cells in bone marrow (BM) within CD45+ leukocytes at the terminal analysis. (B) Frequency of ZsG+ mutant immune cells in PB within CD45+ leukocytes before tumor injection and at the terminal analysis from WT, *Tet2^+/-^* and *Dnmt3a^R^*^878^*^H/+^* CHIP animals. (C) Frequency of ZsG+ mutant immune cells in PB within CD45+CD11b+ myeloid population before tumor injection and at the terminal analysis. (D) Relative enrichment of ZsG+ mutant immune cell frequency in total CD45+ leukocytes, T cells, B cells and myeloid cells in tumors compared to PB from WT young and middle-aged, *Tet2^+/-^*CHIP young and middle-aged and *Dnmt3a^R^*^878^*^H/+^*CHIP young and middle-aged.

**Supplementary Figure 6:**
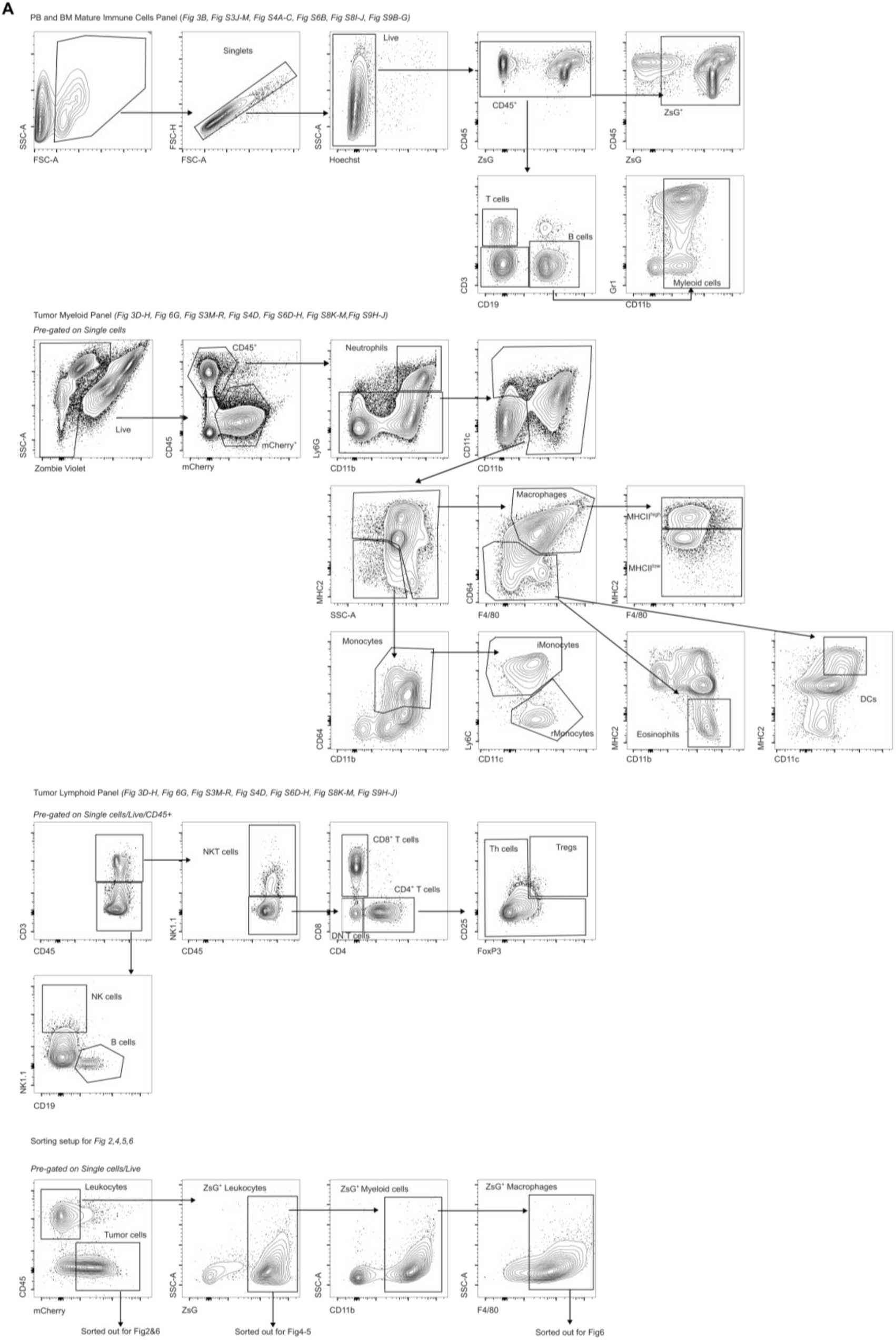
Gating strategies. (A) Gating strategy used in indicated figures.

**Supplementary Figure 7:**
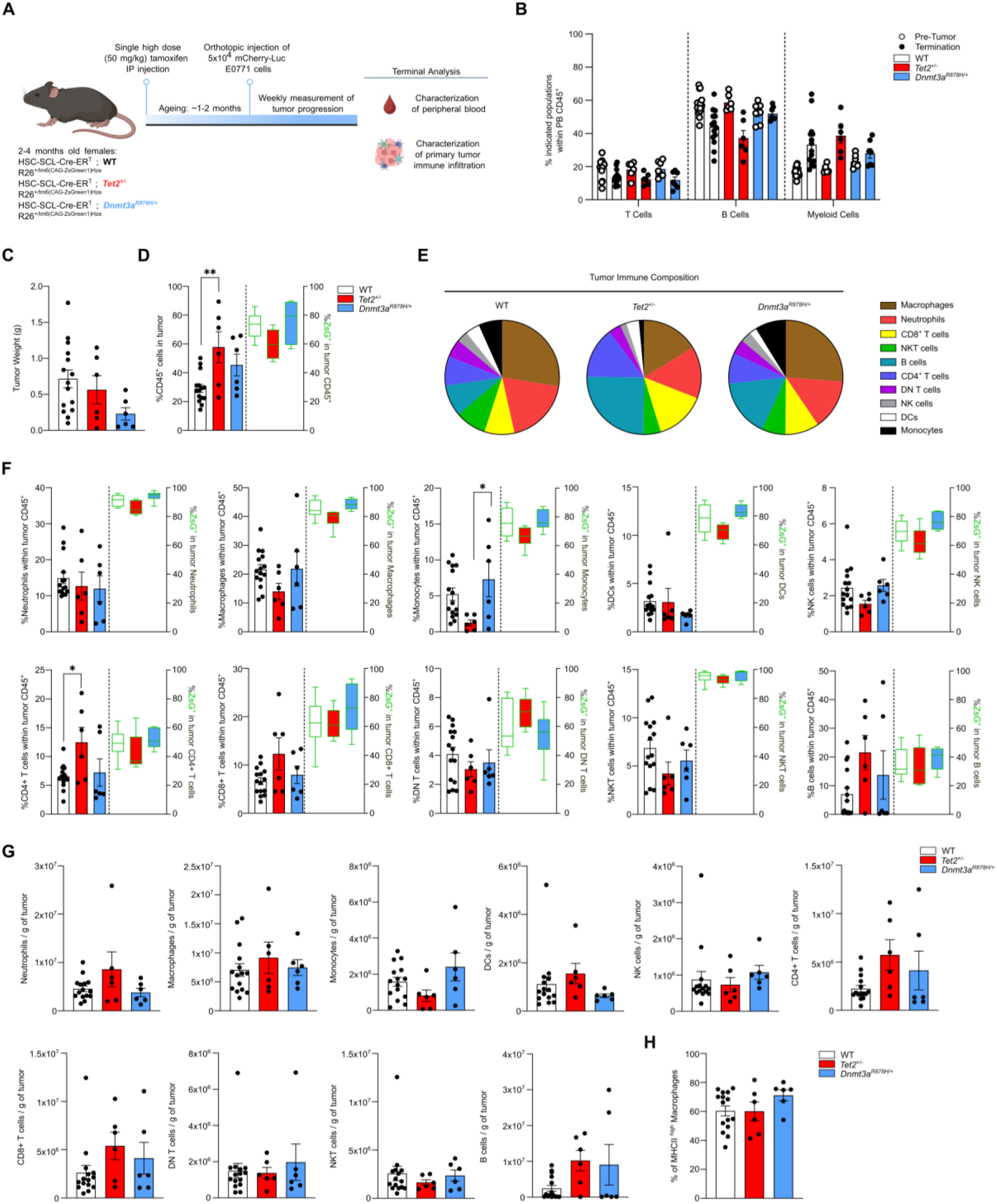
Breast tumors from young mice with *Tet2*-mutant CHIP have similar tumor immune infiltration compared to WT and *Dnmt3a*-mutant CHIP. (A) Graph depicting experimental outline. (B) Frequency of T cells, B cells and myeloid cells in peripheral blood (PB) within CD45+ leukocytes before tumor injection and at the terminal analysis. (C) Tumor weight in grams (g). Data is pooled from the groups shown in Supp. Figure 1 F and M. (D) Percentage of total CD45+ immune cells in total live cells of tumor microenvironment in WT, *Tet2^+/-^* and *Dnmt3a^R^*^878^*^H/+^* CHIP animals by flow cytometric analysis. Frequency of ZsG+ mutant immune cells in the indicated population are depicted on the right y-axis. (E) Pie chart depicting tumor immune composition as a percentage of indicated immune populations in CD45+ leukocytes. (F) Flow cytometric analysis of indicated tumor-infiltrating immune populations from young WT, *Tet2^+/-^* and *Dnmt3a^R^*^878^*^H/+^*CHIP animals. Frequency of ZsG+ mutant immune cells in the indicated population are depicted on the right y-axis. (G) Number of indicated immune cells in the tumor microenvironment normalized to the tumor weight /in grams,g). (H) Fraction of MHC-II^high^ in total macrophages. Statistical analysis was performed by using ordinary one-way ANOVA. Data is shown as mean with SEM. *p<0.05, **p<0.01, ***p<0.001, ****p<0.0001.

**Supplementary Figure 8:**
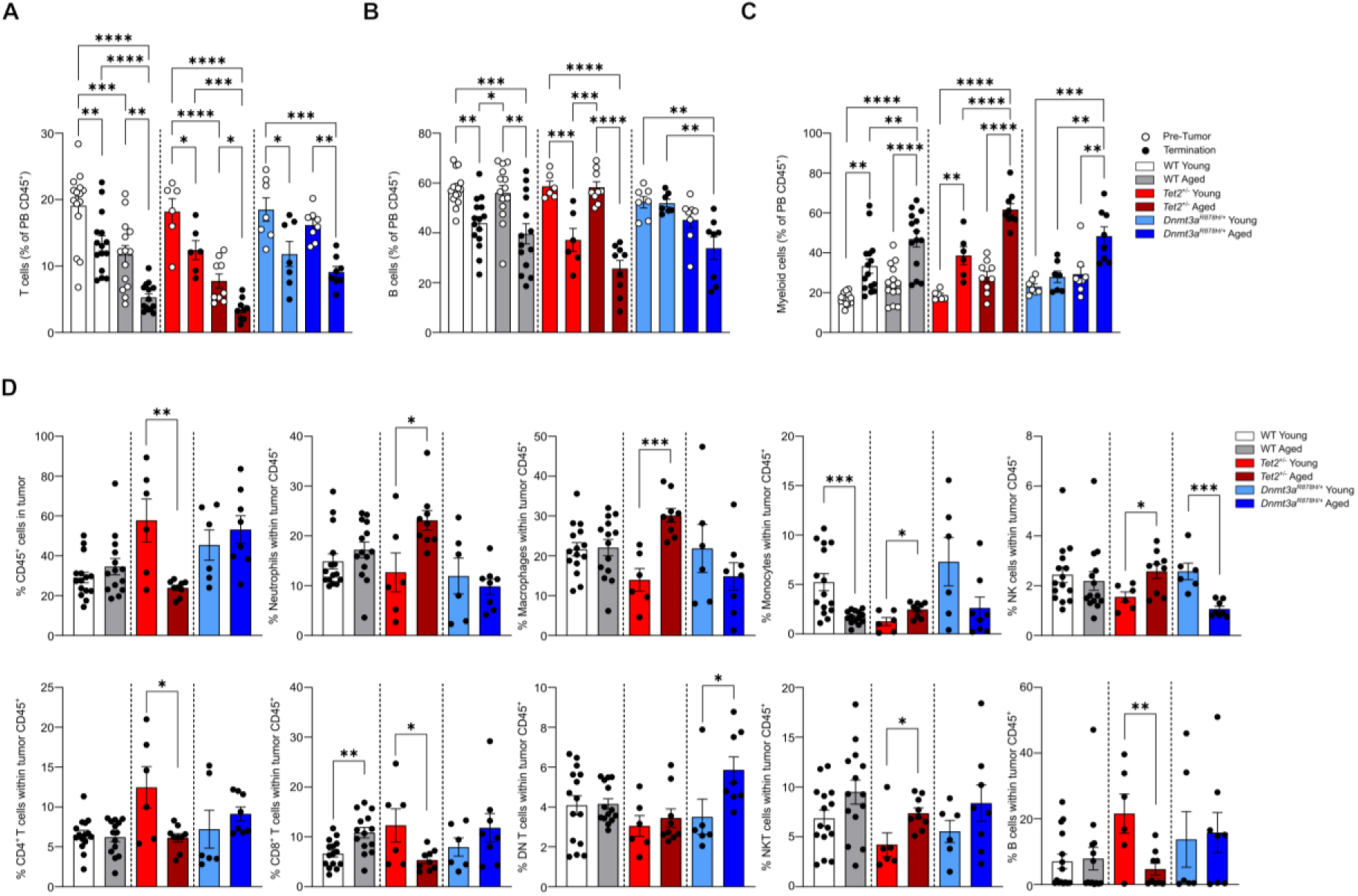
Aging affects the immune composition of breast tumors from *Tet2^+/-^*CHIP mice. (A, B, C) Frequency of T cells, B cells and myeloid cells in peripheral blood (PB) within CD45+ leukocytes before tumor injection and at the terminal analysis in young and middle-aged WT, *Tet2^+/-^* and *Dnmt3a^R^*^878^*^H/+^*CHIP animals. (D) Flow cytometric analysis of indicated tumor-infiltrating immune populations from young and middle-aged WT, *Tet2^+/-^* and *Dnmt3a^R^*^878^*^H/+^*CHIP animals. Data is pooled from the groups shown in Figure 3 and S6.

**Supplementary Figure 9:**
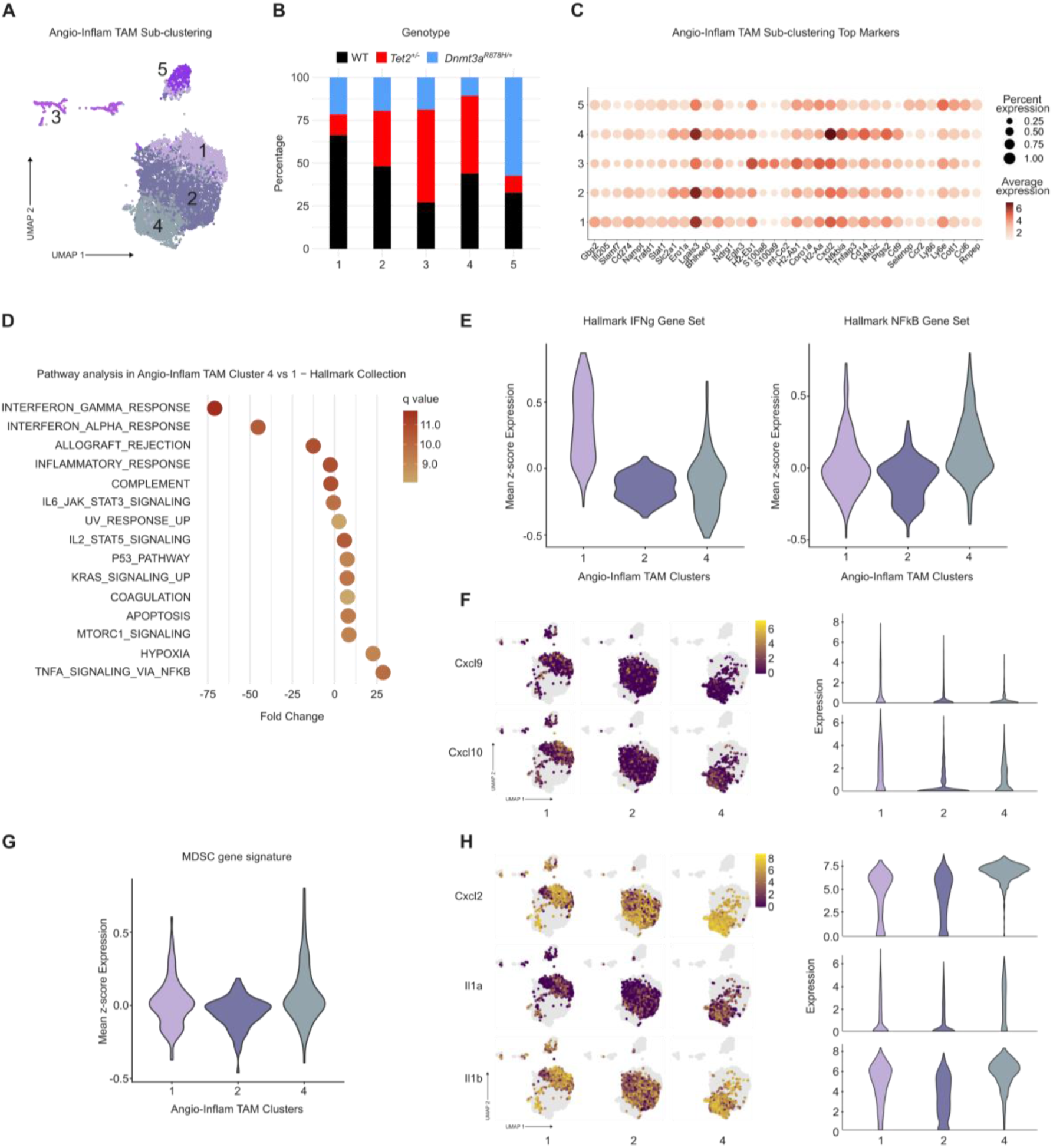
*Tet2*-mutant Angio-Inflammatory TAMs are enriched in clusters with increased NF-kB pathway activation, immunosuppressive signatures and increased IL-1 / Cxcl2 transcription levels compared to WT and *Dnmt3a*-mutant TAMs. (A) UMAP showing the sub-clustering of Angio-Inflammatory TAM population. (B) Distribution of genotypes in each sub-cluster. (C) Dot plot showing the expression of top markers in each sub-cluster of Angio-Inflammatory TAMs. (D) Pathway analysis in Angio-Inflammatory TAM sub-clusters 4 vs 1 in the Hallmark dataset. (E) Violin plots showing the mean z-score for expression of genes in the Hallmark-IFNγ (left) and Hallmark-NFkB (right) gene sets in Angio-Inflammatory TAM sub-clusters. (F) UMAP of sub-clusters of Angio-Inflammatory TAM population showing the expression of Cxcl9 (upper panel), Cxcl10 (lower panel) and respective quantification in violin plots. (G) Violin plots showing the mean z-score for expression of genes in the myeloid-derived suppressor (MDSC) signature. (H) UMAP of sub-clusters of Angio-Inflammatory TAM population showing the expression of Cxcl2 (upper panel), IL-1α (middle panel) and IL-1β (lower panel) and respective quantification in violin plots.

**Supplementary Figure 10:**
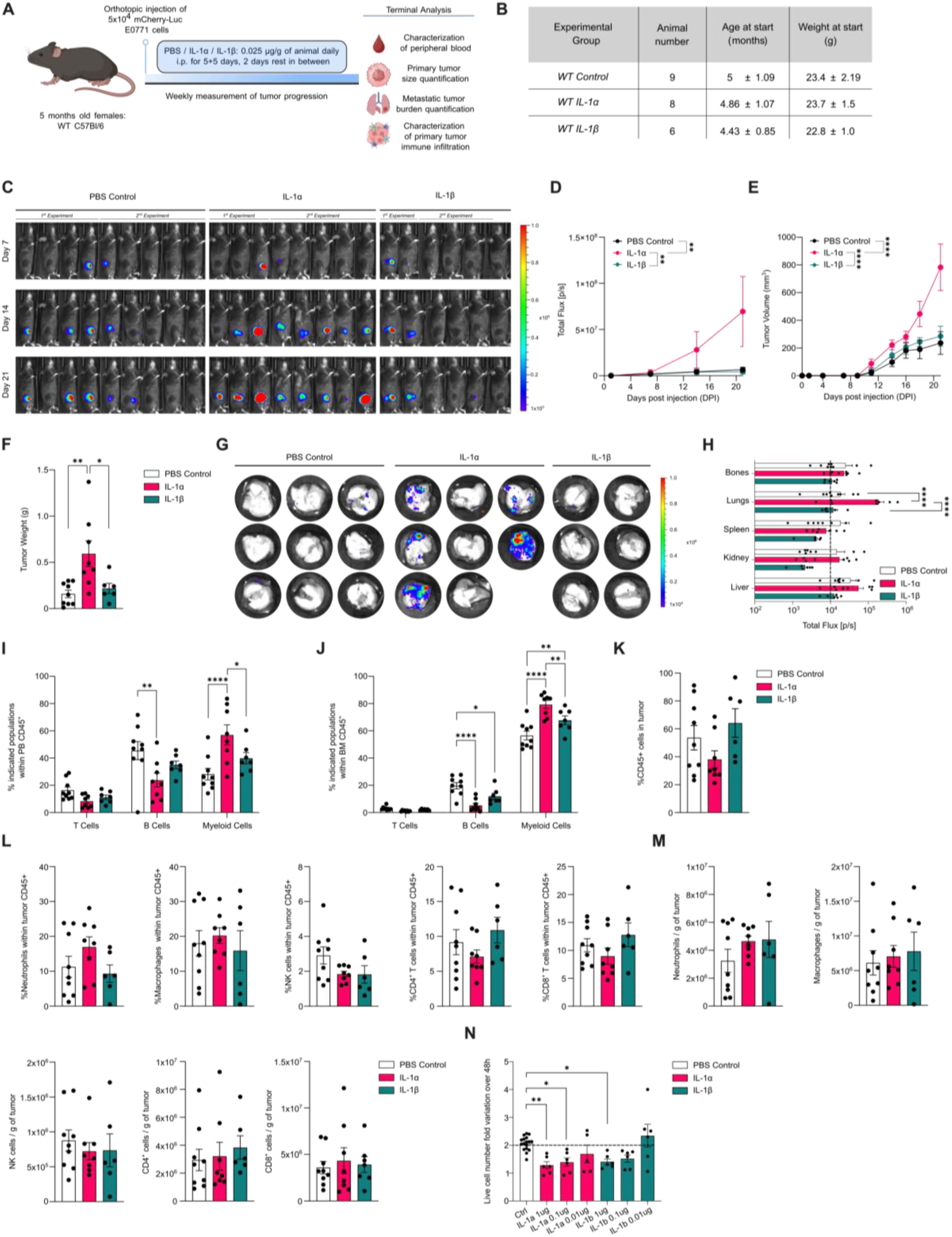
Systemic IL-1α administration promotes breast cancer progression in WT animals. (A) Graph depicting experimental outline. (B) Graph depicting animal numbers, age, weight of each experimental group. (C) *In vivo* growth of orthotopic injected, mCherry-Luciferase expressing E0771 breast cancer over time, evaluated by IVIS imaging of WT animals treated with PBS, IL-1α, and IL-1β. (D) Tumor growth curves (left panel) and individual curves (right panel) by quantification of luminescent intensity from images in (C). (E) Tumor volume measurements over time in mm^3^. (F) Tumor weight measurement in grams at terminal analysis. (G) *Ex vivo* IVIS images of lungs showing level of metastasis in WT animals treated with PBS, IL-1α, and IL-1β. (H) Quantification of luminescent intensity of *ex vivo* images by IVIS showing level of metastasis in indicated organs. Dashed line represents the average background signal (I) Frequency of T cells (CD3+), B cells (CD19+) and myeloid cells (CD3-CD19-CD11b+) in peripheral blood (PB) within CD45+ leukocytes at the terminal analysis. (J) Frequency of T cells, B cells and myeloid cells in bone marrow (BM) within CD45+ leukocytes at the terminal analysis. (K) Fraction of total CD45+ immune cells in total live cells of tumor microenvironment by flow cytometric analysis. (L) Flow cytometric analysis of indicated tumor infiltrating immune populations as a percentage of total CD45+ leukocytes in WT animals treated with PBS, IL-1α, and IL-1β. (M) Number of immune cells in the tumor microenvironment normalized to the gram of tumor for indicated populations. (N) Fold variation of E0771 cell numbers 48 hours after exposure to indicated doses of IL-1α, IL-1β or PBS. For all panels, data is pooled from two independent experiments. For panel N, data is pooled from 3 independent experiments. Statistical analysis was performed by using two-way ANOVA (D, E, H, I, J) and Ordinary one-way ANOVA (F, K, L, M, N). Data is shown as mean with SEM. *p<0.05, **p<0.01, ***p<0.001, ****p<0.0001.

**Supplementary Figure 11:**
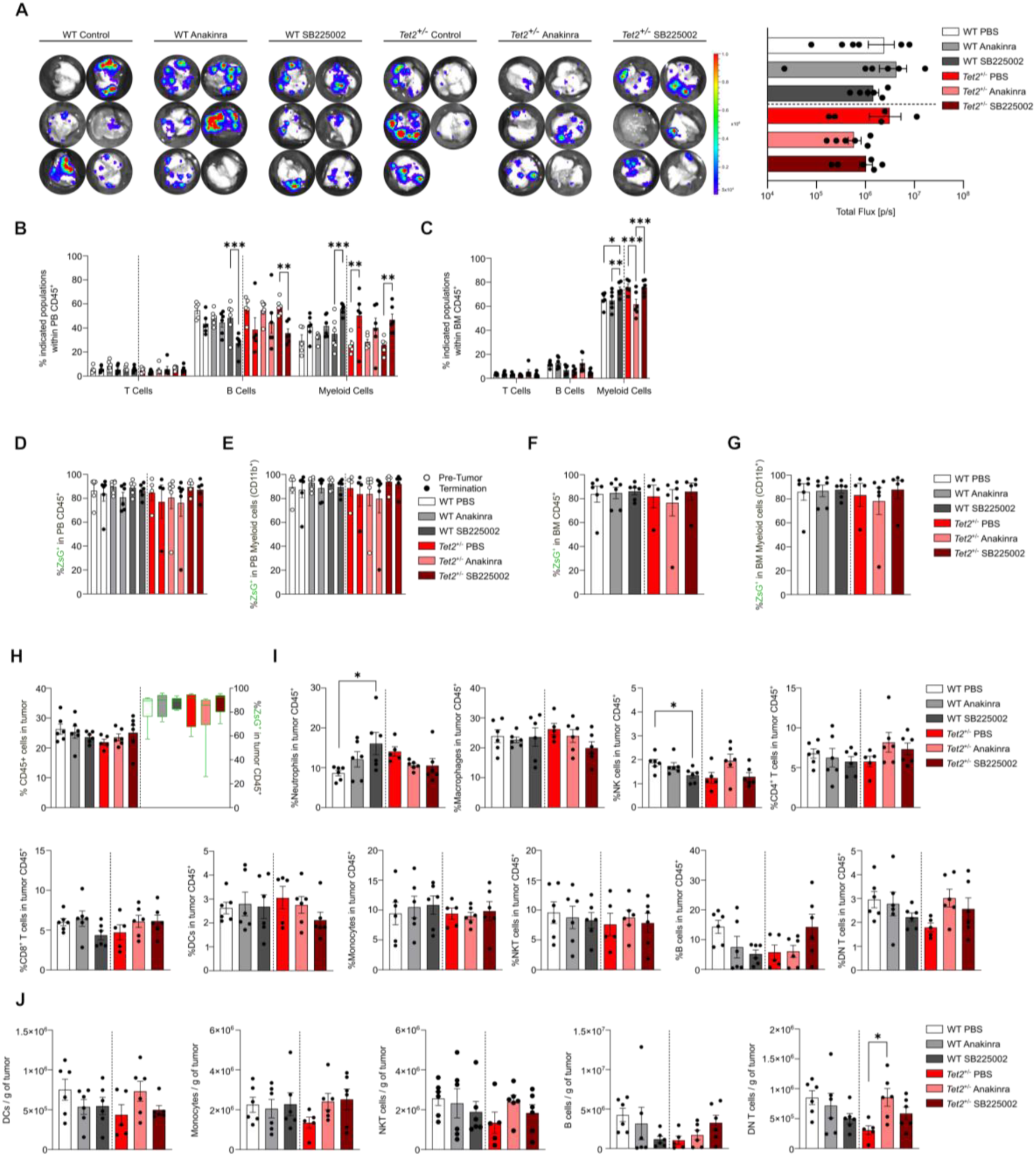
Immune cell characterization in tumors from middle-aged mice with WT or *Tet2*-CHIP treated with IL-1R1 and CXCR2-pathway inhibitors. (A) *Ex vivo* images of lungs by IVIS showing level of metastasis from WT and *Tet2^+/-^*CHIP animals (left panel). Quantification of luminescent intensity of ex vivo images by IVIS showing level of metastasis in lungs (right panel). (B) Frequency of T cells, B cells and myeloid cells in PB within CD45+ leukocytes before tumor injection and at the terminal analysis. (C) Frequency of T cells, B cells and myeloid cells in BM within CD45+ leukocytes at the terminal analysis. (D) Frequency of ZsG+ mutant immune cells in PB within CD45+ leukocytes before tumor injection and at the terminal analysis from WT and *Tet2^+/-^*CHIP animals. (E) Frequency of ZsG+ mutant immune cells in PB within CD45+CD11b+ myeloid population before tumor injection and at the terminal analysis. (F) Frequency of ZsG+ mutant immune cells in BM within CD45+ leukocytes at the terminal analysis. (G) Frequency of ZsG+ mutant immune cells in BM within CD45+CD11b+ myeloid population at the terminal analysis. (H) Percentage of total CD45+ immune cells in total live cells of tumor microenvironment in WT and *Tet2^+/-^*CHIP animals by flow cytometric analysis. Frequency of ZsG+ mutant immune cells in the indicated population are depicted on the right y-axis. (I) Flow cytometric analysis of indicated tumor-infiltrating immune populations from middle-aged WT and *Tet2^+/-^*CH animals with low recombination. Frequency of ZsG+ mutant immune cells in the indicated population are depicted on the right y-axis. (J) Number of indicated immune cells in the tumor microenvironment normalized to the tumor weight /in grams, (g). For all panels, data is pooled from two independent experiments. Statistical analysis was performed by using two-way ANOVA within each genotype (B-C) and ordinary one-way ANOVA within each genotype (remaining panels). Data is shown as mean with SEM. *p<0.05, **p<0.01, ***p<0.001, ****p<0.0001.

## Tables

**Table S1 –** Differential expressed gene (DEG) list observed in E0771 implanted in aged WT, *Tet2^+/-^*-CHIP and *Dnmt3a^fl-R^*^878^*^H/+^*-CHIP mice.

**Table S2 –** Markers of T / NK cells, B cells, dendritic cells, monocytes-macrophages clusters.

**Table S3 –** Markers of T cell sub-clusters.

**Table S4 –** Markers of monocytes-macrophages sub-clusters.

**Table S5 –** Differential expressed gene (DEG) list observed in *Tet2^+/-^*TAMs from aged mice with *Tet2^+/-^*-CHIP after PBS, Anakinra or SB225002 treatment.

## MATERIALS AND METHODS

### Mice

Triple-transgenic mouse strains were generated as described in Caiado et al, 2023. Briefly, HSC-SCL-Cre-ER^T^ ; Tet2^+/flox^; R26^+/tm6^(CAG–ZsGreen1)^Hze^ and HSC-SCL-Cre-ER^T^ ; Tet2^+/+^; R26^+/tm6^(CAG–ZsGreen1)^Hze^, were generated in house by crossing commercially available strains Tet2 floxed (017573 from The Jackson Laboratory), R26^+/tm6^(CAG–ZsGreen1)^Hze^ (007906 from The Jackson Laboratory), and HSC-SCL-Cre-ERt (kindly provided by Radek Skoda). HSC-SCL-Cre-ER^T^; Dnmt3a ^fl-R878H/+^; R26^+/tm6^(CAG–ZsGreen1)^Hze^ were generated in house by crossing commercially available strains Dnmt3a ^tm1Trow^/J (032289 from The Jackson Laboratory), R26^+/tm6^(CAG–ZsGreen1)^Hze^ (007906 from The Jackson Laboratory), and HSC-SCL-Cre-Ert. Wild type C57BL/6 mice were purchased as 2 months old from Charles River Laboratories. Experimental animals were included in experiments when over the lower weight limit for their age group, taking C57BL/6 female mice as standard. All mice were housed and handled according to the guidelines of the Swiss Federal Veterinary Office under specific pathogen-free conditions. All experiments were approved by the Veterinaramt Kanton Zurich (Zurich, Switzerland).

### Cell culture

All cell cultures were performed in 37 ℃ and 5% CO2 in a standard incubator (Binder). E0771 cells (kindly provided by Prof. Maries van den Broek, University of Zurich, Switzerland) were transduced via lentiviral vector to express dual reporter (mCherry-Luciferase). E0771 cells were cultured in DMEM with high glucose with GlutaMAX (Gibco, 10566016) supplemented with 5% fetal bovine serum (FBS, Gibco, A5256801) and 1% penicillin/streptomycin (P/S, Gibco, 15140122). The cells were kept in culture for 5 days before orthotopic injection. Py230 (ATCC, CRL-3279) cells were cultured in Ham’s F12K Medium (Gibco, 21127022) supplemented with 5% FBS, 1% P/S and 0.1% MITO+ serum extender (Corning, 355006). In vitro exposure of E0771 cells to IL-1α or IL-1β was performed by adding 1, 0.1 or 0.01ug/ml of either IL-1α (Recombinant Murine IL-1α, Peprotech, 211-11A) or IL-1β (Recombinant Murine IL-1β, Peprotech, 211-11B) for 48 hours. Cell numbers at 48 hours were normalized to cell numbers at 0 hours.

### In vivo assays

To induce recombination, 2-4 months old female HSC-SCL-Cre-ER^T^ ; Tet2/Dnmt3a^+/+^; R26^+/tm6^(CAG–ZsGreen1)^Hze^ , HSC-SCL-Cre-ER^T^ ; Tet2^+/flox^ ;R26^+/tm6^(CAG–ZsGreen1)^Hze^ and HSC-SCL-Cre-ER^T^ ; Dnmt3a ^fl-R878H/+^; R26^+/tm6^(CAG–ZsGreen1)^Hze^ mice were injected intraperitoneally with high (50 mg/kg) or low (18 mg/kg) doses of tamoxifen. The linear equation for the tamoxifen titration is y= 0.8587x+4.407 where x is the amount of tamoxifen injected (mg/kg of mouse) and y is the expected percentage of ZsG^+^CD45^+^ cells after 5 weeks of injection (Caiado et al, 2023). After tamoxifen injection, mice were allowed to age without any environmental or experimental perturbations. Measurement of peripheral blood chimerism before tumor inoculation was done by collecting peripheral blood in 50-75 µl from sublingual veins. For orthotopic breast tumor model, 5#x00D7;10^4^ E0771 cells were injected into the 4^th^ inguinal mammary fat pad of the respective animals. Tumor progression over time was measured using a caliper and tumor volume is calculated using the formula: V= 0.5 x L x W^2^ where L is the longer diameter and W is the shorter diameter of the tumor. Tumor progression was also measured via bioluminescence imaging using In Vivo Imaging System (IVIS) (PerkinElmer). Mice received an intraperitoneal injection of 150 mg/kg D-luciferin bioluminescent substrate (15 mg/ml stock solution in PBS, XenoLight, PerkinElmer, 122799) and allowed for a 5-minute period. Mice were anesthetized under 2% of isoflurane and imaging was carried out for 2 minutes exposure time with the default settings. Bioluminescent intensity was quantified using total flux (photons/seconds) via Living Image software (version 4.7.1). For antagonist treatments, 7 days after tumor inoculation, Anakinra (Kineret, Sobi) and SB225002 (MedChemExpress, HY-16711) were intraperitoneally administered with a dose of 500 µg/mouse every other day and 20 mg/kg for 2 rounds of 5 consecutive days with 2-day rest in between respectively. In vivo tumor progression was measured as described above. After 7 days of tumor inoculation, 5 months old wild type C57Bl/6 females were administered intraperitoneally with PBS, IL-1α (Recombinant Murine IL-1 α, Peprotech, 211-11A) or IL-1β (Recombinant Murine IL-1β, Peprotech, 211-11B) with a dose of 0.025 µg / g of mouse in phosphate buffered saline (PBS) with 1% bovine serum albumin (BSA) for 2 rounds of 5 consecutive days with 2 day rest in between.

### Isolation of bone marrow and peripheral blood cells

Bone marrow (BM) cells were harvested from femurs and tibias. Single cell suspensions were prepared by flushing bones in FACS buffer (2%FBS, 2mM EDTA in PBS). Red blood cells were lysed with 5 ml red blood cell lysis buffer (150 mM NH4Cl, 10 mM KHCO3, and 0.1 mM Na2EDTA) for 5 minutes and cells were washed in 5 ml FACS buffer. Cell debris was removed with 70-μm filters (BD). For peripheral blood, red blood cells were lysed with 1ml of red cell lysis buffer for 10 minutes and cells were washed once with 2 ml FACS buffer.

### Ex vivo organ imaging

Mice were injected intraperitoneally with 15 mg/kg D-luciferin. After 5 minutes, animals were euthanized and tumors, livers, spleens, lungs, kidneys, both femurs and tibias immediately collected and imaged using IVIS system with 5 minutes of exposure. Bioluminescent intensity was quantified using total flux (photons/seconds) via Living Image software.

### Tumor and Lung Dissociation

Tumors and lungs were harvested into PBS supplemented with 2% FBS. Single cell suspension was generated by enzymatic and mechanical digestion using dissociation medium as previously described (de la Fuente et al, 2021, StemCellTechnologies protocol,2021). 5 ml medium was prepared for each tissue by combining 3.75 mL DMEM+GlutaMAX with 500 µL of a 50x collagenase/hyaluronidase stock (final concentrations: 0.365 mg/mL collagenase and 0.125 mg/mL hyaluronidase, Collagenase from Clostridium histolyticum, Sigma, C9891, Hyaluronidase from bovine testes, Sigma, H3506) and 750 µL DNase I from 1 mg/mL stock solution in PBS (final concentration: 0.15 mg/mL, Roche, 7002221). Tissues were incubated at 37 ℃ for 25 minutes. After enzymatic dissociation, tissues were strained with 70 µm cell strainer (Falcon, 352350) and red blood cells were lysed with 5 ml of red blood cell lysis buffer for 5 minutes and cells were washed once with 10 ml FACS buffer.

### Flow Cytometry

All staining protocols were performed using FACS buffer. Cells were incubated with respective antibodies for 30 minutes at 4℃ (see Supplementary Table). Live/dead cell selection was performed using two different viability dyes. Hoechst 33342 (Thermo Fisher Scientific,62249) was added to PB and BM cells 5 minutes prior to acquisition following the respective antibody staining. Tumor cells were incubated with ZombieViolet (BioLegend, 423113) for 20 minutes at 4℃ prior to antibody labelling. To block the unspecific binding, tumor cells were incubated with TruStain FcX (anti-mouse CD16/32) antibody (BioLegend, 101319) for 10 minutes at room temperature. Intracellular staining of Foxp3 in the lymphoid panel was performed using Foxp3 Transcription Factor Staining Buffer Set (eBioscience, 00-5523-00). Counting beads (Sigma-Aldrich, P7458) were used for absolute counts. Stained cells were acquired using BD LSR Fortessa II flow cytometer and analyzed with FlowJo software (v10.10.0). All the immune populations were gated on Live/Singlets/CD45+. In PB analysis, T cells are defined by the CD3 expression, B cells with CD19 and myeloid cells with CD11b expression. In tumor myeloid analysis, markers of each population as follows: neutrophils (CD11b^+^ Ly6G^+^), macrophages (Ly6G^-^CD11b^+^ CD11c^+^ MHCII^+^ F4/80^+^ CD64^+^), monocytes (Ly6G^-^CD11b^+^ CD11c^+^ MHCII^-^CD64^+^), dendritic cells (DC) (Ly6G^-^CD11b^+^ CD11c^+^ MHCII^+^ F4/80^-^CD64^-^) (PMID: 26938654). In tumor lymphoid analysis, markers of each population as follows: NKT cells (CD3+ NK1.1+), CD8+ T cells (CD3+ NK1.1-CD8+), CD4+ T cells (CD3+ NK1.1-CD4+), NK cells (CD3-NK1.1+) and B cells (CD3-CD19+) (PMID: 34261667).

### Histology

Lungs were harvested into 4% paraformaldehyde (PFA) in PBS and stored overnight at 4°C. Tissues were then routinely embedded in paraffin, and 5-µm-thick sections were stained with hematoxylin and eosin (HE) at the Institute for Pathology and Molecular Pathology at University Hospital Zurich. The stained slides were digitalized using a NanoZoomer slide scanner (Hamamatsu Photonics, Hamamatsu City, Japan), with 40× objective lens setup. All lung sections were examined by a board-certified veterinary pathologist using QuPath (v0.7.0), and metastatic lesions were manually annotated and enumerated.

### Single -cell RNA Sequencing

Cells isolated from tumors were sorted into two different populations: ZsG^+^CD45^+^ *Tet*-mutant leukocytes and mCherry^+^ E0771 tumor cells (see Supplementary Figure 5 for gating strategy) using BD Aria III 4L. 1-2#x00D7;10^6^ ZsG^+^CD45^+^ cells were sorted into FACS buffer and proceeded for fixation following the protocol (Fixation of Cells & Nuclei for Chromium Fixed RNA Profiling, 10X Genomics). Single-cell RNA sequencing was performed using 10X Genomics Chromium System (10x Genomics, CA). cDNA libraries were generated according to the manufacturer’s instructions (Single Cell Gene Expression Flex) and sequenced using Illumina NovaSeq X Plus (Paired End 150 bp) at Functional Genomics Center Zurich (FGCZ).

### Single-cell RNA Sequencing Data Processing

Raw sequencing data were processed with Cell Ranger^69^ via the FGCZ online platform SUSHI, generating filtered gene-cell matrices for downstream analysis in R using the SingleCellExperiment package. Quality control was performed using the following settings: cells expressing more than 5% mitochondrial genes, a number of detected genes outside the 500-8,000 range, or unique UMIs outside 1,000-100,000 were excluded. Normalization and variance decomposition were performed per sample. The results of multiple variance decomposition were combined using combineVar function from scran. The samples were combined using multiBatchNorm from the batchelor R package. PCA and UMAP were performed using top 20% of highly variable genes (HVGs). We used the scDblFinder to detect and remove doublets. Cells were clustered using k-means and graph-based clustering. Markers of each cluster were identified using the scoreMarkers function from the scran R package. Next, clusters were next annotated based on marker genes and previous studies to identify immune cell subpopulations. Monocytes/macrophages and T/NK cells were next subsetted and the analysis was performed as described above starting with identifying top 20% HVGs and rerunning PCA, UMAP and clustering. This allowed further annotated monocytes/macrophages and T cells sub-populations. Specific annotated monocyte/macrophage subpopulations were then further sub-clustered to identify cellular stated linked with the genotypes in our study. We next applied pathway analysis to gain functional insights into those cellular states. Pathway analysis as a two-group comparison was performed using the SCPA R package and the MSigDB gene sets. Cells from each subpopulation were down sampled to 1000 to reduce the computational cost.

We selected the following immune populations for the cellular interaction analysis: cytotoxic T cells, NK cells, neutrophils and all the monocyte/macrophage clusters excluding unconventional monocytes. The cellular interaction analysis was performed using CellChat and CellChatDB for mouse. The significant cell-cell interactions were determined using CellChat default parameters, with population size accounted for and ‘‘trimean’’ used for calculating mean gene expressions for the *Dnmt3a-* and *Tet2*-mutant selected immune populations. Subsequently, the CellChat vignette for comparison across different biological conditions was used with default settings.

### Bulk RNA Sequencing

Total RNA extraction was performed by following the protocol of RNeasy Plus Mini Kit (Qiagen, 74134). 2#x00D7;10^6^ mCherry^+^ E0771 tumor cells and 2-5#x00D7;10^5^ macrophages (CD11b^+^ F4/80^+^) were sorted into FACS buffer with BD Aria III 4L. Dried cell pellets were used to extract total RNA content. Bulk RNA sequencing was performed using High Throughput Next Generation Sequencing system. cDNA libraries were generated according to the manufacturer’s instructions using Tecan Universal Plus mRNA-Seq protocol and sequenced using Illumina NovaSeq X Plus (Paired End 150 bp) at Functional Genomics Center Zurich (FGCZ). Quality control and mapping using Kallisto software was performed using the SUSHI bioinformatics pipeline. Exploratory data analysis and differential expressions were performed using DESeq2 R package according to the DESeq2 vignette. GSEA analysis was run with ClusterProfiler R package. The results were visualized using the following packages: EnhancedVolcano, pheatmap, ggplot and ComplexUpset.

### Statistical Analysis

Statistical analysis was performed using GraphPad Prism (v11.0.01). For statistical tests for each panel, see the respective figure legends.

